# ATR coordinates nuclear and cytoskeletal mechanics during leukemic migration

**DOI:** 10.64898/2026.11.17.767255

**Authors:** Marina Armesto, Natalia Bustos, Edurne Armesto-Longueira, Ivett Barany, Gracia Peralta-Carrero, Ana Benito, Elena G. Sánchez, Manuel Ramírez, Ignacio Benedicto, Javier Redondo-Muñoz

## Abstract

Mechanical adaptation is essential for tumor cells to cross physical barriers during dissemination, but the mechanisms linking genome-stress signalling to cellular mechanics remain poorly understood. Here, we demonstrate that ATR (ataxia telangiectasia and Rad3-related kinase) acts as a dual regulator of nuclear and cytoskeletal adaptations in T-cell acute lymphoblastic leukemia (T-ALL). Genetic depletion or pharmacological inhibition of ATR altered nuclear architecture and mechanical behaviour, increasing nuclear deformability in 3D confined conditions. Mechanistically, ATR inhibition increased chromatin accessibility, H3K4me3 levels and the association of WDR5 with chromatin. In parallel, ATR interference reduced myosin light chain kinase (MLCK) expression, ERM phosphorylation and the association between ERM proteins and actin, consistent with altered cortical cytoskeletal regulation. These changes had distinct consequences in leukemic cell migration: ATR inhibition promoted T-ALL migration through confined 3D environments but impaired transendothelial migration and reduced early homing to peripheral tissues in vivo. Inhibition of Chk-1 did not reproduce the nuclear response to ATR inhibition, suggesting that these effects might not depend on the ATR-Chk-1 axis signalling. Together, our findings identify ATR as a regulator of nuclear and cortical organization during leukemic migration, revealing that enhanced nuclear deformability might be insufficient to promote migration across diverse physical microenvironments.

## INTRODUCTION

Nuclear mechanics and mechanosensing integrate extracellular forces to nuclear organization and gene regulation, controlling multiple cell functions including genome integrity, cell migration, and differentiation (1,2). During tumor dissemination, cells must adapt to distinct mechanical and adhesive environments, including confined extracellular spaces and endothelial barriers, which impose different requirements for cancer cell migration (3,4). Mechanical adaptation to these environments requires coordinated changes in the nucleus and the cytoskeleton that allow cells to deform and move through confined regions (5,6). However, nuclear deformation during migration can also damage chromatin, compromising genome integrity and cell viability (7–10).

Ataxia telangiectasia and Rad3-related kinase (ATR) is a central regulator of replication stress signalling and genome stability, coordinating DNA replication, checkpoint activation, and repair processes in response to cellular stress (11,12). Beyond its canonical role in the DNA damage response (13), ATR contributes to cellular responses to mechanical stress and to the maintenance of nuclear integrity during cell migration (14,15). ATR inhibition has also been shown to sensitize cancer cells to cell death induced by genotoxic treatments, including acute lymphoblastic leukemia (ALL) cells (16–19). ALL is the most common childhood cancer, and T-cell ALL (T-ALL) is a subset of ALL associated with a substantial risk of relapse and treatment failure (20). Our previous work showed that epigenetic changes contribute to nuclear deformability and the invasive behaviour of ALL cells in vitro and in vivo (21,22). However, how ATR may link genome-stress signalling to nuclear adaptation during leukemic cell migration remains unclear. At the same time, successful cell migration requires not only nuclear deformability, but also coordinated adhesion, cortical cytoskeletal organisation and force generation to cross the endothelium. Whether ATR regulates these distinct mechanical requirements in a coordinated manner has not been established.

Here, we investigated the role of ATR in the mechanical adaptation of T-ALL cells by examining nuclear architecture, chromatin organization, cytoskeletal regulation and leukemic cell migration in different environments. We found that genetic or pharmacological targeting of ATR altered nuclear mechanical behaviour and promoted migration through confined 3D environments, but reduced transendothelial migration *in vitro* and early leukemic cell recovery from peripheral tissues *in vivo*. These effects were accompanied by increased H3K4me3 levels and WDR5 association with chromatin. Pharmacological inhibition of WDR5, a core subunit involved in H3K4 methylation, attenuated the nuclear deformation and 3D invasion induced by ATR inhibition, suggesting that WDR5 contributes to these responses induced by ATR. Targeting ATR also altered cortical cytoskeletal regulation, including reduced myosin light chain kinase (MLCK) expression, ERM (ezrin/radixin/moesin) phosphorylation and ERM-actin interactions. Finally, inhibition of Chk-1 did not reproduce the nuclear response to ATR inhibition. Together, these findings suggest that ATR coordinates distinct mechanical functions during leukemic migration and that increased nuclear deformability alone is insufficient for efficient leukemic dissemination through different physical barriers.

## EXPERIMENTAL SECTION

### Cell culture

Jurkat (RRID: CVCL_0065) and CCRF-CEM (RRID: CVCL_0207) cells were obtained from ATCC. Primary human umbilical vein endothelial cells HUVECs were obtained from Lonza and cultured in Endothelial Cell Growth Basal Medium (EBM) supplemented with EGM SingleQuots. Primary samples from pediatric patients with ALL, all under 16 years of age, were obtained with informed consent under protocols approved by the Institutional Review Board of the Niño Jesús University Hospital and CSIC. Patients were diagnosed and treated according to the SEHOP-PETHEMA 2013 protocol (Spanish Society of Pediatric Hematology and Oncology – Spanish Program for the Treatment of Hematologic Diseases), and patient characteristics are detailed in **Table 1**. ALL cell lines and primary ALL cells were maintained in RPMI 1640 supplemented with 10% FBS, 1.8 mM L-glutamine and 25 mM HEPES. All cells were maintained at 37 °C and 5% CO₂. Cell lines were routinely tested for mycoplasma contamination. Cells were treated for 24 h with VE-821 (1 µM for cell lines; 10 µM for primary samples), PF-477736 (250 nM) or OICR-9429 (1 µM). ML-7 was used at 20 µM for 20 min.

### ATR silencing

ATR (sc-29763-SH) or control (sc-108060) were purchased from Santa Cruz Biotechnology. Jurkat cells (1 × 10⁶) were seeded in 800 µL of RPMI 1640 and transfected with 2.5 µL of shRNA plasmid solution (0.25 µg) and 7.5 µL of UltraCruz® Transfection Reagent, according to manufacturer’s instructions. Non-targeting shRNA plasmid was used as control conditions. Cells were collected 24 h after transfection, and ATR silencing efficiency was verified by Western Blot before experiments.

### Immunofluorescence

Cells under different conditions were seeded onto 10 µg/mL poly-L-lysine-coated glass slides for 30 min (1 h for cell spreading analysis), fixed in 4% paraformaldehyde (10 min at room temperature, RT), permeabilized with 0.5% Triton X-100 in PBS (5 min), blocked in PBS containing 10% FBS and 0.1% Triton X-100 (30 min, RT) and incubated with appropriate primary antibodies (1 h, RT). After several washes, samples were incubated with secondary antibodies (1 h, RT) and Alexa Fluor 568-conjugated phalloidin to stain F-actin (1 h, RT). After nuclear staining with Hoechst 33342 (10 min, RT), samples were washed with PBS and water, and mounted with DABCO mounting medium. Images were acquired on a Leica DMi8 inverted confocal microscope using an HC PL APO 63×/1.30 NA glycerol immersion objective. Image quantification and analysis were performed with ImageJ/Fiji (National Institutes of Health).

### Nuclear isolation

Cells under different conditions were resuspended in Buffer A (10 mM HEPES, 10 mM KCl, 1.5 mM MgCl₂, 0.34 M sucrose, 10% (v/v) glycerol, 1 mM DTT, 0.1% Triton X-100, and protease inhibitor cocktail) for 5 min on ice. Following centrifugation at 1,360 x g (5 min, 4 °C), the pellet with isolated nuclei were resuspended in PBS or TKMC (50 mM Tris [pH 7.5], 25 mM KCl, 3 mM MgCl₂, 3 mM CaCl₂, and protease inhibitor cocktail) buffers for subsequent analyses.

### Nuclear shrinking

Isolated nuclei from different conditions were resuspended in TKMC buffer and sedimented onto poly-L-lysine-coated plates (10 µg/mL). Then, nuclei were incubated or not with additional 5 mM MgCl₂ (15 min), fixed, permeabilized, stained with Hoechst 33342, and mounted with DABCO mounting medium on a slide. Images were acquired on a Leica DMi8 inverted confocal microscope using an HC PL APO 63×/1.30 NA glycerol immersion objective. Quantification and analysis of the nuclear area were performed with ImageJ/Fiji.

### Atomic Force Microscopy (AFM)

8 × 10⁵ isolated nuclei from different conditions were resuspended in PBS and seeded on 50 mm cell culture dishes (Willco GWST-5040) coated with 100 µg/mL poly-L-lysine following the protocol (8,9). AFM measurements were performed with a Bruker Bioscope Resolve mounted on an inverted microscope (Nikon Eclipse Ti2) connected to an ORCA-Flash4.0LT (Hamamatsu) camera. Indentation curves were fitted within Nanoscope Analysis (Bruker) using a cone-sphere model. Imaging was performed in filtered PBS at RT. Pre-calibrated Silicon Tip – Nitride cantilevers (PFQNM-LC-V2 Bruker) were used with a 70 nm tip radius. PeakForce Tapping/PeakForce Capture with a maximum indentation force of 0.5 nN was used.

### Mechanical compression

Isolated nuclei were resuspended in PBS, stained with Hoechst 33342 and sedimented onto 10 µg/mL poly-L-lysine-coated plates and placed in the CSOW 110 1-well cell confiner device (4Dcell). Following the manufacturer’s instructions, mechanical confinement was performed by pushing the nuclei with a glass slide containing 3 µm-high micropillars. Images of nuclei were taken with a HC PL APO 63×/1.30 NA glycerol immersion objective before and after the confinement, and analysis of the nuclear area was performed with ImageJ/Fiji.

### Western blot

Cells under different conditions were lysed in Laemmli Buffer, sonicated at 75% amplitude for a total of 5 min (30 s ON/30 s OFF in cycles) using a using a Q500 sonicator (Qsonica), boiled at 95 °C for 5 min, and loaded onto SDS-polyacrylamide gels (7.5–15%). Proteins separated by electrophoresis were transferred to nitrocellulose membranes, blocked in 3% milk in TBS-Tween (1 h) at RT and incubated overnight with primary antibodies at 4 °C. Membranes were washed in TBS-Tween, incubated with appropriate HRP-conjugated secondary antibodies, and protein signals were detected in a ChemiDoc MP Imaging System (Bio-Rad). For the stripping process, 3 stripping buffer washes (1% Glycine [pH 2.2], 1% SDS and 0.05% NP-40) of 30 min were performed before blocking the membranes and reprobing with specific antibodies. For cellular fractionation, the cytoplasmic fraction was extracted in buffer A (5 min, 4 °C) and centrifuged for 5 min at 4 °C at 3,500 x g. Nuclear pellets were resuspended in RIPA buffer (50 mM Tris-HCl [pH 7.4], 134 mM NaCl, 0.1% SDS, 1% NP-40, 0.5% Sodium deoxycholate) containing protease inhibitors. After 15 min, soluble nuclear fractions were collected from centrifugation (5 min, 4 °C, 17,200 x g) and chromatin bound fractions were resuspended in Laemmli Buffer and sonicated. Proteins were resolved by SDS-PAGE. Quantification and analysis of images were performed using ImageJ/Fiji software.

### Cell invasion and nuclear morphology in 3D conditions

Collagen gels were prepared inside Transwell inserts at a final concentration of 1.7 mg/mL bovine collagen type I neutralized with 0.1 volume of a solution of 7.5% NaHCO₃ and 25 mM HEPES in RPMI (1 h, 37 °C). 5 × 10⁵ cells in 100 µL of serum-free RPMI were seeded onto the collagen gel and RPMI medium with 10% FBS was added to the bottom chamber. After 24 h, collagen gels were fixed with 4% PFA (30 min at 37 °C, 30 min at RT, and 30 min at 4 °C), permeabilized with 0.5% Triton X-100 (30 min) and stained with Propidium Iodide (overnight, 4 °C). Invading cells were imaged by capturing confocal z planes every 10 µm on a Leica DMi8 inverted confocal microscope with a HC ACS APO 10×/NA 0.3 dry objective. 3D reconstructions were performed with LAS-AF 3D software from Leica and the percentage of penetrating cells was quantified with the ImageJ/Fiji software. To visualize the nuclear morphology, cells were embedded in collagen gels and allowed to migrate randomly. Then, cells were fixed, permeabilized, and their nuclei were stained with Hoechst 33342. Images of the gels were captured on a Leica DMi8 inverted confocal microscope using a HC ACS APO 40×/NA 1.3 glycerol immersion objective. Nuclear circularity was analyzed using the shape descriptor analysis tool on the ImageJ/Fiji software.

### Statistical Analysis

Statistical analyses were performed using GraphPad Prism 8. Independent cell-culture experiments, individual patient samples and individual animals constituted biological replicates, as applicable. Each experiment included at least two biological replicates. The numbers of cells or nuclei analysed (*n*) and the numbers of independent replicates (*N*) are reported separately in the figure legends. Data are presented as mean ± SD unless otherwise indicated. Comparisons between two groups were performed using two-sided unpaired or paired parametric or non-parametric t-tests (Student’s t-test, Mann–Whitney U test, or Wilcoxon signed-rank test), as appropriate for the experimental design and data distribution. Comparisons involving three or more groups were performed using one and two-way ANOVA followed by Tukey’s multiple-comparisons test. Statistical significance was indicated as follows: (*) P<0.05; (**) P<0.01; (***) P<0.001. Statistical methods and multiple-testing corrections for RNA-seq, ChIP-seq and gene set enrichment analyses are described in the corresponding Experimental subsections.

## RESULTS

### ATR interference remodels nuclear architecture and increases nuclear deformation under mechanical compression

ATR has been implicated in the maintenance of nuclear integrity and in the response of cells to mechanical stress (23). To determine whether ATR contributes to nuclear adaptation in T-ALL cells, we first analyzed nuclear morphology in Jurkat and CCRF-CEM cells cultured on poly-L-lysine. Pharmacological inhibition of ATR with VE-821 increased the nuclear area compared to control cells (**Figure 1A, 1B and Suppl. Figure S1A, S1B**), without significant changes observed in cell cycle or viability at the time points analyzed (**Suppl. Figure S1C, S1D**). Genetic depletion of ATR using shRNA (**Suppl. Figure S1E**) produced a similar increase in nuclear area (**Figure 1C, 1D**), supporting that the nuclear phenotype was associated with ATR depletion rather than other off target effects of VE-821. Given that primary tumor cells typically exhibit higher resistance than immortalized cell lines (24), we optimized VE-821 concentration to ensure effective ATR inhibition and to analyze its effect in primary T-ALL samples. VE-821 treatment also increased nuclear area by 10.3% to 49.7% in primary cells from all three different T-ALL patients (**Suppl. Figure S1F, S1G**), indicating that ATR inhibition also alters nuclear morphology in primary leukemic cells. Because changes in nuclear morphology can be associated with alterations in chromatin organization (25), we next analyzed chromatin accessibility using DNase I sensitivity. Jurkat and CCRF-CEM cells treated with VE-821 showed increased DNase I sensitivity compared to control cells, consistent with a more accessible chromatin state after ATR inhibition (**Figure 1E and Suppl. Figure S1H**).

**Figure 1.**
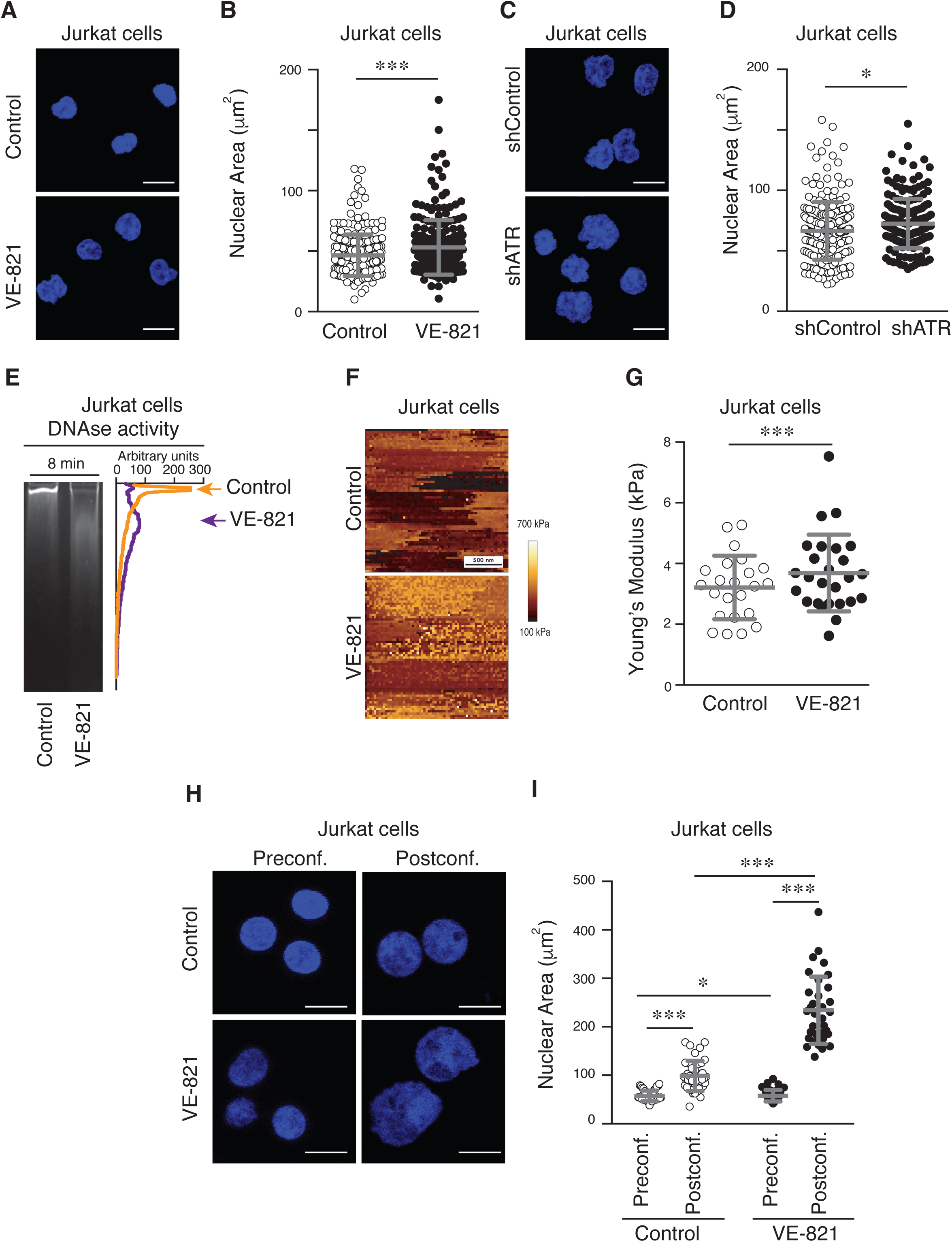
ATR interference alters nuclear morphology and nuclear biomechanical response in T-ALL cells. **(A)** Jurkat cells were treated with or without VE-821 (1 µM for 24 h). Nuclear morphology was then visualized by Hoechst staining (DNA) (scale bar: 10 µm). **(B)** Nuclear area quantification (*n* = 228-235 cells; *N* = 3). *** P<0.001. **(C)** Nuclear morphology of control (shControl) and ATR-depleted (shATR) Jurkat cells visualized by Hoechst staining (scale bar: 10 µm). **(D)** Nuclear area quantification (*n* = 234-195 cells; *N* = 3). * P<0.05. **(E)** Representative gel image and corresponding relative-intensity profile of control and VE-821-treated Jurkat cells following treatment with DNase I for 15 min. Arrowheads indicate the highest DNA-digested peak. **(F)** Isolated nuclei from control and VE-821-treated Jurkat cells were seeded on poly-L-lysine-coated dishes and imaged by AFM (atomic force microscopy). PeakForce Tapping image of a representative nucleus captured with a maximum indentation force of 0.5 nN. **(G)** Elastic modulus measurements using AFM of control and VE-821-treated Jurkat cells. (*n* = 23-26 nuclei; *N* = 3). *** P<0.001. **(H)** Representative images of control and VE-821-treated Jurkat cells under mechanical compression visualized by Hoechst staining (scale bar: 10 µm). **(I)** Nuclear areaquantification before and after mechanical compression (*n* = 234/98-190/65 nuclei; *N* = 2). *** P<0.001.

We next asked whether these changes in nuclear morphology were accompanied by alterations in nuclear mechanical behaviour. First, we added Mg^2+^ to isolated nuclei from control and VE-821-treated cells to determine how the nucleus might respond to shrinking conditions. We observed a reduction of the 8% of nuclear area in control conditions upon Mg^2+^ addition, with a greater reduction of the 53% in the nuclear area of VE-821-treated cells after Mg^2+^ addition (**Suppl. Figure S2A, S2B)**. To further investigate mechanical properties, we analyzed isolated nuclei by atomic force microscopy (AFM) and found that VE-821 treatment increased the surface heterogeneity of the nucleus and the Young’s modulus (**Figure 1F, 1G and Suppl. Figure S2C**). This result is consistent with previous observations that changes in the Young’s modulus measures local topography of the nucleus rather than the integrated biomechanical response of the nucleus (26). We therefore analyzed the deformation of isolated nuclei under mechanical compression to determine how ATR-targeted nuclei respond to external mechanical forces. We found that isolated nuclei from ATR-targeted cells showed greater change in the nuclear area than those from control cells, indicating an altered mechanical response to confinement (**Figure 1H, 1I and Suppl. Figure S2D, S2E**). Together with the changes in nuclear morphology and AFM measurements, these results indicate that ATR inhibition alters the mechanical behaviour of the T-ALL nucleus and increases its capacity to adapt to confined environments.

### Targeting ATR promotes nuclear deformation and migration through confined 3D environments

Nuclear deformability is an important constraint for cell migration through confined environments (27). Since ATR inhibition altered nuclear mechanical behaviour, we next analyzed whether this phenotype affected leukemic cell migration. We cultured control and VE-821-treated T-ALL cells on top of collagen matrices and analyzed their capacity to invade through the 3D environment in response to serum, keeping an additional condition of control cells in the absence of any stimulus as a negative control of cell migration (**Figure 2A**). In contrast to the reduced migration previously reported after ATR targeting in other cellular models (13), VE-821 treatment increased the invasion of both T-ALL cell lines through confined 3D conditions (**Figure 2B and Suppl. Figure S3A and S3B**). We also confirmed in primary T-ALL cells that VE-821 treatment increased the percentage of invasive cells into the collagen gel (**Suppl. Figure S3C and S3D**). Similarly, ATR depletion modestly increased the invasive capacity of Jurkat cells (**Figure 2C and 2D**). ATR primarily signals through Chk-1 during the canonical replication stress response (33); however, increasing evidence indicates that ATR and Chk-1 have independent functions (34). We therefore asked whether the nuclear and migratory phenotypes observed after ATR inhibition could be explained by inhibition of this canonical ATR–Chk-1 pathway. First, we found that treating T-ALL cells with a selective Chk-1 inhibitor (PF-477736) did not increase T-ALL cell invasion into 3D collagen gels (**Suppl. Figure S3E-H**). We next analyzed nuclear morphology of embedded cells migrating randomly through collagen matrices. ATR-targeted cells showed reduced nuclear circularity in collagen, consistent with greater nuclear deformation under these conditions (**Figure 2E-2H and Suppl. Figure S4A and S4B**). This effect was also reproducible in primary cells from T-ALL patients (**Suppl. Figure S4C**). Moreover, we found that unlike VE-821 treatment, Chk1 inhibition did not reduce the nuclear circularity of Jurkat cells in 3D conditions, and even increased the circularity of CCRF-CEM cells (**Suppl. Figure S4D-G**). Together, these results indicate that ATR modulates the capacity of T-ALL cells to adapt their nuclei and migrate through confined 3D environments via a mechanism that is at least partly separable from its canonical Chk-1-dependent replication stress response.

**Figure 2.**
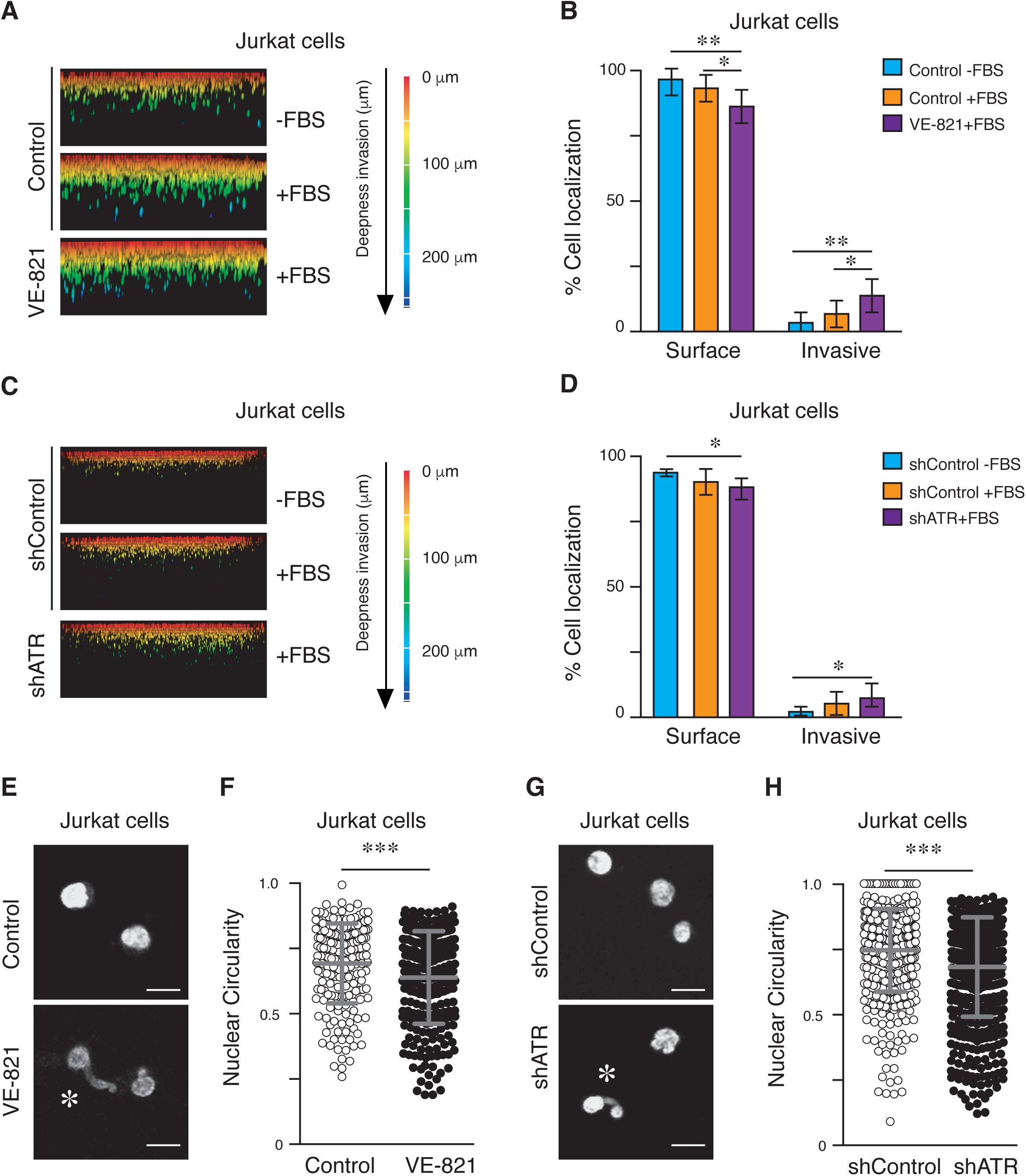
Targeting ATR enhances nuclear deformation and migration of T-ALL cells through confined 3D environments. **(A)** Representative 3D reconstructions of control and VE-821-treated Jurkat cells migrating towards FBS through a 1.7 mg/mL collagen I matrix for 24 h. Cells were fixed and stained with Propidium Iodide. Colours indicate invasion depth (scale bar: 250 µm). **(B)** Quantification of the percentage of total cell-occupied area in the collagen gel. We designated the upper third as “Surface cells” (non-invasive), while cells in the lower two-thirds within the collagen were designated as “Invasive cells” (*N* = 3). * P<0.05, ** P<0.01. **(C)** Representative 3D reconstructions of control (shControl) and ATR-depleted (shATR) Jurkat cells migrating through a collagen I matrix as in (A). Colours indicate invasion depth (scale bar: 250 µm). **(D)** Quantification of the percentage of total cell-occupied area in the gel. (*N* = 3). * P<0.05. **(E)** Representative immunofluorescence images of control and VE-821-treated Jurkat cells embedded in a 1.7 mg/mL collagen I matrix for 4 h, fixed and stained with Hoechst. Asterisks indicate high deformable nuclei (scale bar: 10 µm). **(F)** Nuclear circularity index quantification (*n* = 199-255 cells; *N* = 3). *** P<0.001. **(G)** Representative immunofluorescence images of control and ATR-depleted Jurkat cells embedded in a collagen I matrix, fixed, and stained with Hoechst. Asterisks indicate highly deformable nuclei (scale bar: 10 µm). **(H)** Nuclear circularity index quantification (*n* = 471-521 cells; *N* = 3). *** P<0.001.

### ATR interference increases H3K4me3 levels and WDR5 association with chromatin

Our previous work established that WDR5-dependent H3K4 methylation contributes to chromatin remodeling and nuclear mechanical adaptation during invasion through confined 3D environments (32). Since ATR inhibition also increased the cell migration through 3D conditions, we hypothesized that there might be a connection between ATR and H3K4 methylation. We found that VE-821 treatment increased the bulk levels of H3K4me3 (**Figure 3A, 3B and Suppl. Figure S5A**). ATR-depleted cells also showed increased levels of H3K4me3 (**Figure 3C, 3D**); however, we confirmed that Chk-1 inhibition did not affect this epigenetic marker (**Figure 3E**, **Figure 3F and Suppl Figure S5B**), suggesting that the nuclear response observed after ATR inhibition was not reproduced by Chk-1 inhibition. To investigate how ATR regulates H3K4 methylation, we performed biochemical fractionation of cytosol, nuclear and chromatin-bound proteins and observed that the distribution of WDR5 and other components of the H3K4 methylation machinery, such as ASH2L, were enriched in the chromatin fraction of cells treated with VE-821 (**Figure 3G**). To examine whether WDR5 contributed to the nuclear and invasive responses, we combined VE-821 with the WDR5 inhibitor OICR-9429. Blocking WDR5 activity abrogated the decrease in nuclear circularity induced by VE-821 (**Suppl. Figure S5C and S5D**). Moreover, WDR5 inhibition attenuated the cell invasion effect induced by VE-821 (**Suppl. Figure S5E**). Together, our data suggest a functional link between cell invasion and nuclear circularity induced by targeting ATR and the H3K4 methylation and WDR5.

**Figure 3.**
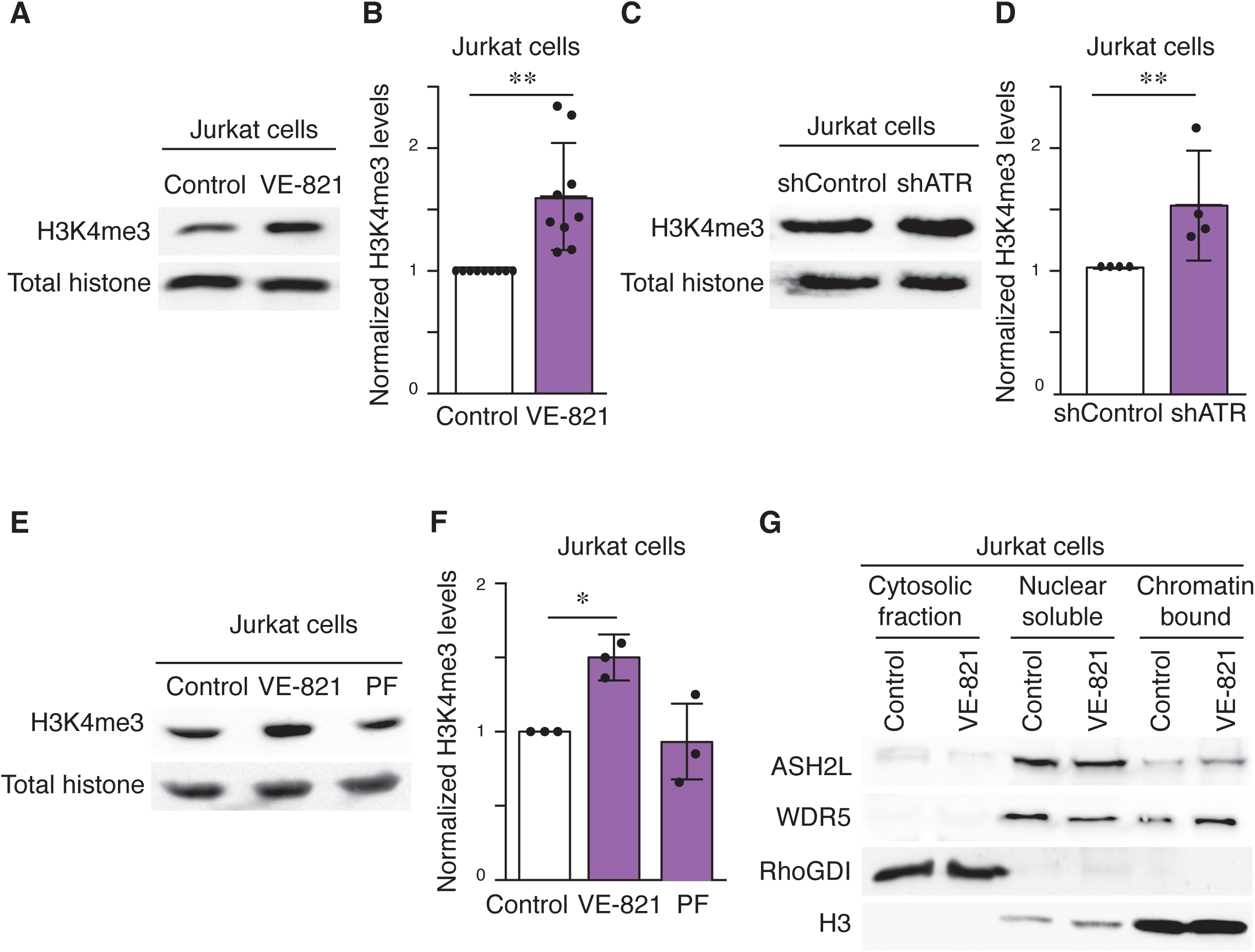
ATR interference increases H3K4me3 levels in T-ALL cells. **(A)** Representative immunoblots showing H3K4me3 levels in total lysates from control and VE-821-treated Jurkat cells. **(B)** H3K4me3 / Total histone ratio. (*N* = 9). ** P<0.01. **(C)** Representative immunoblots showing H3K4me3 levels in total lysates from control (shControl) and ATR-depleted (shATR) Jurkat cells. **(D)** H3K4me3 / Total histone ratio. (*N* = 4). ** P<0.01. **(E)** Representative immunoblots showing H3K4me3 levels in total lysates from control and Chk-1-inhibited (250 nM PF-477736 for 24 h) Jurkat cells. **(F)** H3K4me3 / Total histone ratio. (*N* = 3). * P<0.05 **(G)** Representative immunoblots showing WDR5 and ASH2L protein levels in cytoplasmic, nuclear, and chromatin-bound fractions from control and VE-821-treated Jurkat cells. RhoGDI and Histone H3 were used as internal control of cytoplasmic and chromatin fractions, respectively.

### Integrated multi-omic analysis reveals transcriptional program and H3K4me3 chromatin remodeling upon ATR targeting

Given the increase in global H3K4me3 levels, we next examined the genomic distribution of H3K4me3 by ChIP-seq. Following library-size normalization, H3K4me3 profiles were broadly similar between control and VE-821-treated cells, with no widespread increase in normalized signal intensity (**Figure 4A, Suppl. Figure S6A**). H3K4me3 peaks were predominantly located in proximal promoter regions in both cell lines (**Suppl. Figure S6B and S6C**), consistent with their enrichment near Transcription Start Sites (TSS). UpSet analysis further delineated the intersection among promoter, exonic, intronic, and untranslated-region annotations assigned to these peaks (**Suppl. Figure S6D**). Gene Ontology (GO) enrichment analysis of the H3K4me3 peaks identified categories related to chromatin and transcriptional regulation in CCRF-CEM cells, and transcriptional regulation, actin binding and kinase-regulatory functions in Jurkat cells (**Figure 4B**). These analyses described distinct functional compositions of the H3K4me3-associated gene sets across T-ALL cell lines.

**Figure 4.**
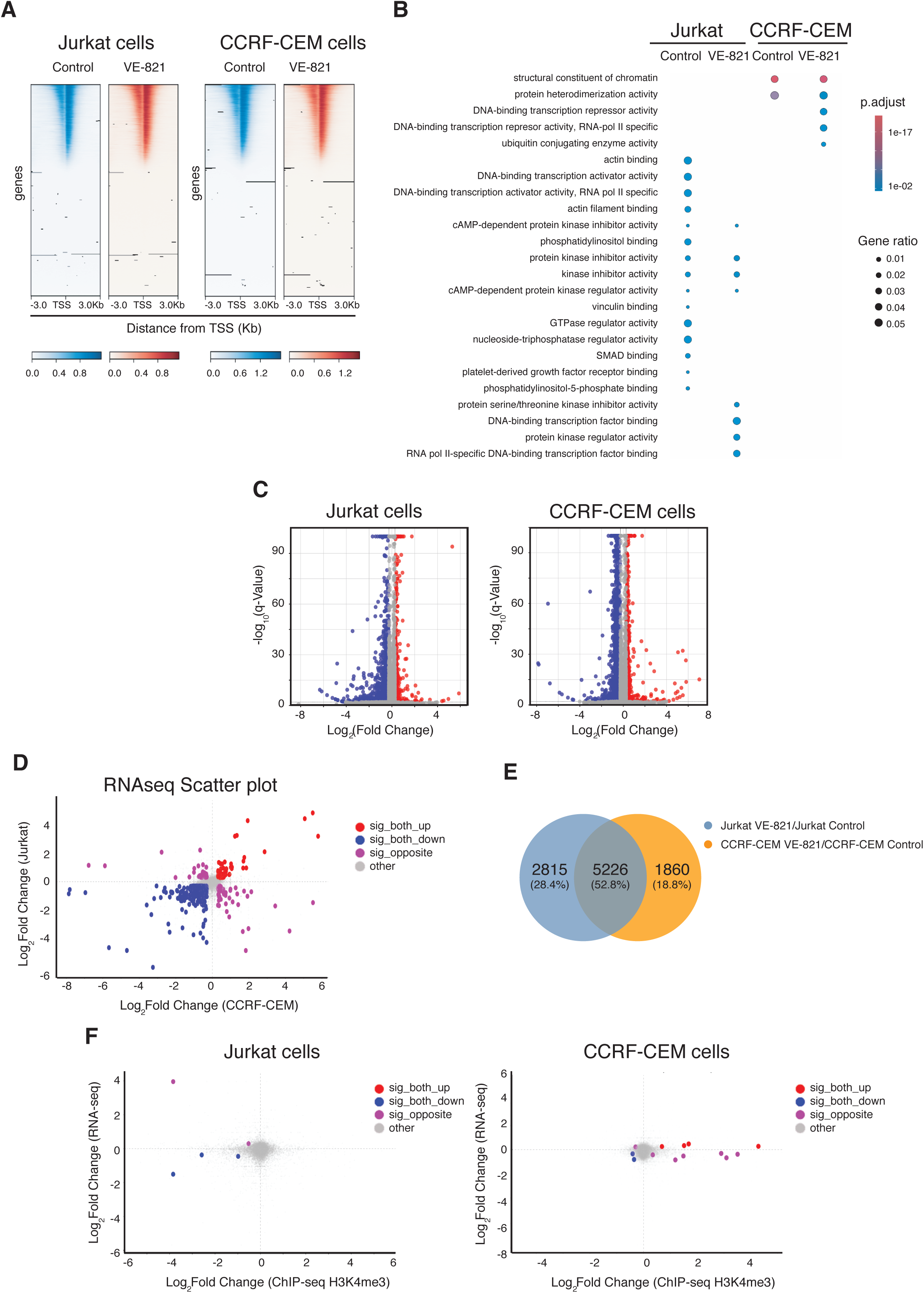
ATR inhibition reshapes transcriptional programs and promotes H3K4me3-associated chromatin remodeling in T-ALL cells. **(A)** H3K4me3 signal distribution plots centered around the Transcription Start Sites (TSS; ±3.0 kb) across genome-wide gene regions in control and VE-821-treated Jurkat cells (left) and CCRF-CEM cells (right). **(B)** Dot plot of Gene Ontology (GO) enrichment analysis showing significantly enriched Molecular Function terms across the different cell lines and treatment groups. Dot size represents the Gene Ratio, and the color scale reflects the adjusted significance (p.adjust value). **(C)** Volcano plots illustrate transcriptomic changes in control and VE-821-treated Jurkat cells (left) and CCRF-CEM cells (right). The x-axis represents the Log₂(Fold Change), and the y-axis shows the -Log_10_(q-Value). Significantly upregulated genes are highlighted in red, and downregulated genes are in blue. **(D)** Scatter plot comparing the Log₂(Fold Change) values of VE-821 treatment in Jurkat cells (y-axis) versus CCRF-CEM cells (x-axis). Dots are color-coded to indicate genes significantly upregulated in both cell lines (red), downregulated in both (blue), oppositely regulated (purple), or non-significant/other (grey). **(E)** Venn diagram illustrating the unique and shared differentially expressed genes (DEGs) between VE-821-treated vs. control in Jurkat cells (green) and in CCRF-CEM cells (orange). **(F)** Scatter plots for Jurkat cells (left) and CCRF-CEM cells (right) comparing gene-associated Log₂(Fold Change) of ChIP-seq H3K4me3, x-axis and Log₂(Fold Change) of RNA-seq, y-axis.

To examine the gene expression response to ATR inhibition, we performed RNA-seq in Jurkat and CCRF-CEM cells treated with VE-821. Differential expression analysis identified changes in steady-state mRNA abundance in both cell lines (**Figure 4C and Suppl. Figure S7A)**. Venn Diagram showed a substantial gene overlapping between cell lines (**Figure 4D**); a comparative scatter plot of log₂ fold changes identified genes in both lines showed concordant and opposite responses between the cell lines (**Figure 4E**), suggesting that ATR inhibition might induce different cell-line-dependent changes. Gene Set Enrichment Analysis (GSEA) identified positive enrichment of ribosomal components and mitochondrial inner membrane complex gene sets in both cell lines. Specifically, Jurkat cells showed negatively enriched categories related to stimulus detection and selected receptor activities (**Suppl. Figure S7B**), whereas CCRF-CEM cells exhibited enrichment of gene sets associated with extracellular structure organization, collagen metabolism and extracellular matrix components (**Suppl. Figure S7C**). These results indicate that the response to ATR inhibition includes shared mitochondrial and ribosomal signatures, whilst other gene sets differ between the two cell lines. Our transcriptional analysis identified that Jurkat cells showed reduced expression of several actomyosin- and cytoskeletal organization-associated genes, such as *MYLK* gene that encodes for the protein MLCK, whereas CCRF-CEM cells exhibited a distinct response involving both upregulated and downregulated cytoskeletal regulators. Notably, the convergence of actin-binding signatures in Jurkat cells together with transcriptional changes involving extracellular matrix/collagen regulation in CCRF-CEM cells points to a potential effect of ATR on cytoskeletal organization and cell–environment interactions.

Finally, to assess the relationship between H3K4me3 and transcriptional responses, we integrated promoter-associated H3K4me3 signals with the corresponding RNA-seq changes in each cell line. The integrated plots revealed heterogeneous relationships between the two measurements, indicating that differences in H3K4me3 signal were not consistently accompanied by changes in mRNA abundance (**Figure 4F**, **Table 2**). Together, these results show that ATR inhibition alters gene expression landscape in T-ALL cells, without a uniform relationship between estimated changes in H3K4me3 signal and mRNA abundance. These findings highlight the broad impact of ATR inhibition on cellular regulatory programs and suggest alterations in pathways governing receptor-associated adhesion and cellular motility that may have functional consequences for leukemic cell behavior and dissemination.

### ATR interference alters early leukemic tissue homing and transendothelial migration

Tumor cells face additional adhesive and physical barriers during dissemination, including vascular adhesion and transendothelial migration (28). To determine whether the observed mechanical, transcriptional, and epigenetic alterations affect leukemic cell dynamics and their ability to interact with vascular barriers and infiltrate peripheral tissues, we next investigated early leukemic dissemination in vivo upon ATR inhibition. We injected control and VE-821-treated T-ALL cells intravenously into immunocompromised mice and, 3 h after injection, quantified leukemic cells recovered from peripheral tissues. For both cell lines the treatment with VE-821 reduced the number of leukemic cells present in the bone marrow, showing further inhibitory effects into other organs for CCRF-CEM cells (**Figure 5A and 5B**). A possible explanation for the altered organ distribution of ATR-targeted cells is defective transendothelial migration despite their higher invasiveness in 3D conditions. Therefore, we next performed an in vitro transendothelial migration assay using HUVEC monolayers. We found that ATR inhibition also significantly reduced the ability of cells to cross the endothelial barrier (**Figure 5C and 5D)**. Thus, ATR inhibition produced opposite effects on two migration settings: it increased invasion through confined 3D matrices but reduced transendothelial migration and in vivo homing of leukemic cells into the bone marrow. Because processes like tissue homing and transendothelial migration requires the coordinated regulation of surface receptors and cortical cytoskeleton dynamics (29), and our multi-omics data point us on this direction, we explored whether ATR interference impairs the mechanical and adhesive machinery, such as receptor expression, actin polymerization, and cortical tension, required for leukemic cells to cross endothelial barriers and colonize peripheral tissues. First, we observed that VE-821 treatment did not substantially alter the expression of integrins, chemokine receptors, or total F-actin levels (**Suppl. Figure S8A-D**), suggesting that the reduced transendothelial migration was not explained by a general loss of adhesion receptors or actin abundance. Moreover, we found that ATR interference did not alter the static adhesion to VCAM-1 in both T-ALL cell lines (**Figure 5E and 5F**).

**Figure 5.**
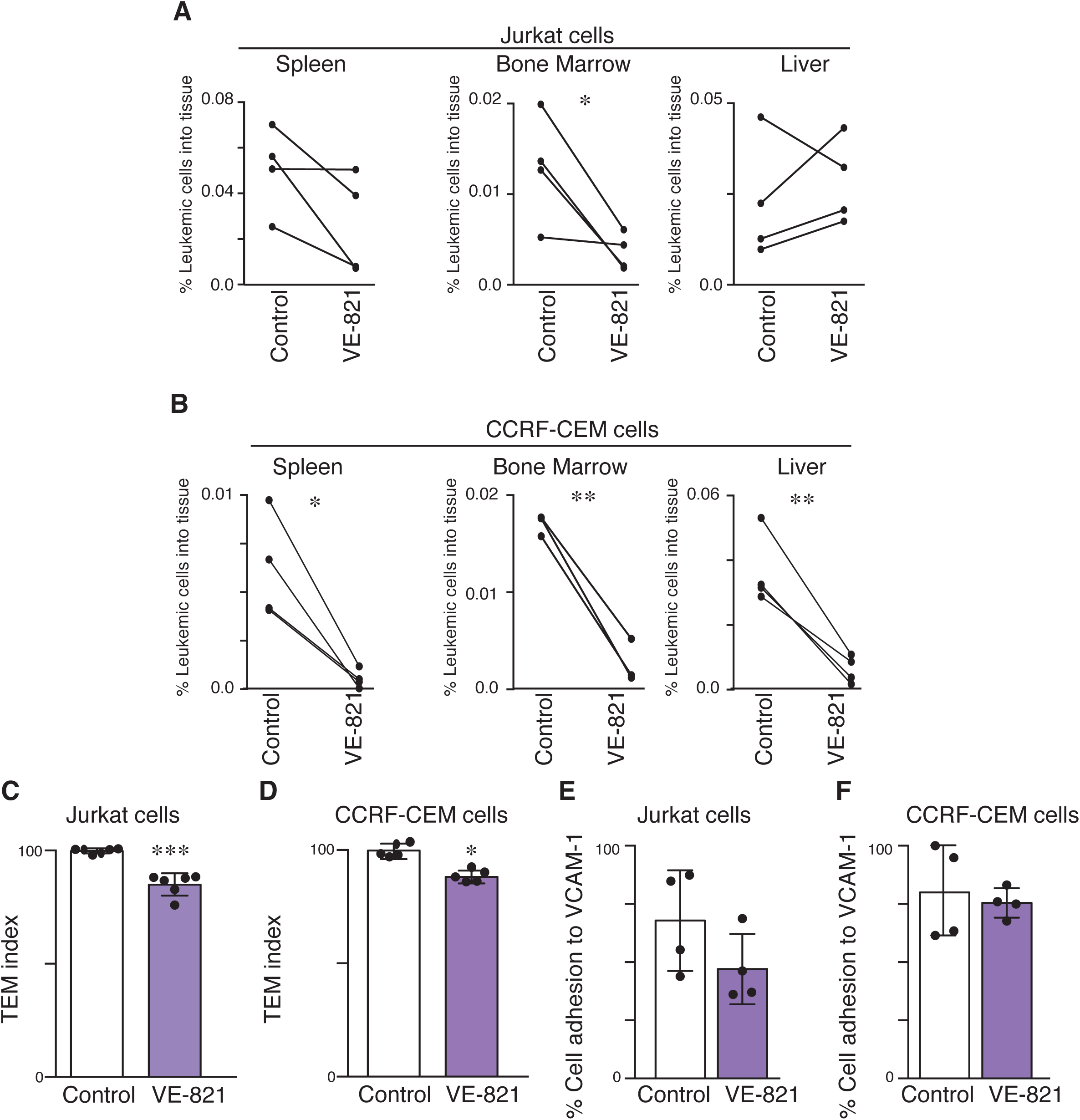
ATR inhibition impairs early leukemic tissue dissemination and transendothelial migration. **(A)** Control and VE-821-treated Jurkat cells were labelled with living dyes, mixed at a 1:1 ratio, and injected intravenously into NOD scid gamma (NSG) mice. After 3 h, the distribution of labelled cells in the bone marrow, spleen, and liver was assessed by flow cytometry. The percentage of labelled leukemic cells recovered from each tissue was determined relative to the total number of events (*N* = 4 mice). * P<0.05. **(B)** Control and VE-821-treated CCRF-CEM cells were labelled, mixed, and injected intravenously into NSG mice as in (A). After 3 h, the distribution of labelled cells was assessed by flow cytometry. (*N* = 3). * P<0.05, ** P<0.01. **(C)** Control and VE-821-treated Jurkat cells were labelled with living dyes, mixed at a 1:1 ratio, and seeded onto a HUVEC monolayer. Migration across the endothelial barrier and through 5-µm-pore Transwell filters was quantified. The graph shows the ratio of migrated cells (*N* = 6). *** P<0.001. **(D)** Transendothelial migration through a HUVEC monolayer of control and VE-821-treated CCRF-CEM cells was quantified as in (C). (*N* = 5). * P<0.05. **(E)** Quantification of cell adhesion to VCAM-1-coated coverslips in control and VE-821-treated Jurkat cells (*N* = 4). **(F)** Quantification of cell adhesion to VCAM-1-coated coverslips in control and VE-821-treated CCRF-CEM cells (*N* = 4).

Next, we focused on determining how ATR interference might be compromising the cortical actomyosin cytoskeleton, which is a key determinant of the transendothelial migration and invasion of T-ALL cells. First, we studied the cellular spreading and found that T-ALL spreading on 2D substrates was increased after VE-821 treatment compared with control cells (**Figure 6A, 6B, and Suppl. Figure S9A and S9B**). We next analyzed proteins involved in cortical actomyosin regulation, such as MLCK and ERM proteins, which contribute to coupling cortical actin to the plasma membrane and regulate nuclear changes (30,31). ATR interference reduced MLCK and phosphorylated ERM protein levels compared with control cells (**Figure 6C and Suppl. Figure S9C**), suggesting altered machinery controlling actomyosin contractility. Interestingly, we found that the treatment with PF-477736 recapitulated the effects of VE-821 in the phosphorylation status of ERM proteins (**Figure 6D**). As a complementary approach, we treated T-ALL cells with the MLCK inhibitor ML-7. Consistent with VE-821 treatment, ML-7 increased cell spreading (**Suppl. Figure S9D-9G**). To further test ERM associations with the actin cytoskeleton, we performed co-immunoprecipitation experiments. Under the extraction conditions used, ATR inhibition reduced the recovery of actin in ERM immunoprecipitates (**Figure 3E**), and the ERM binding in the reciprocal immunoprecipitation of actin (**Figure 3F and Suppl. Figure S9H**). These results indicate that ATR inhibition alters the regulation of the cortical cytoskeleton, which might contribute to the reduced transendothelial migration observed after targeting ATR.

**Figure 6.**
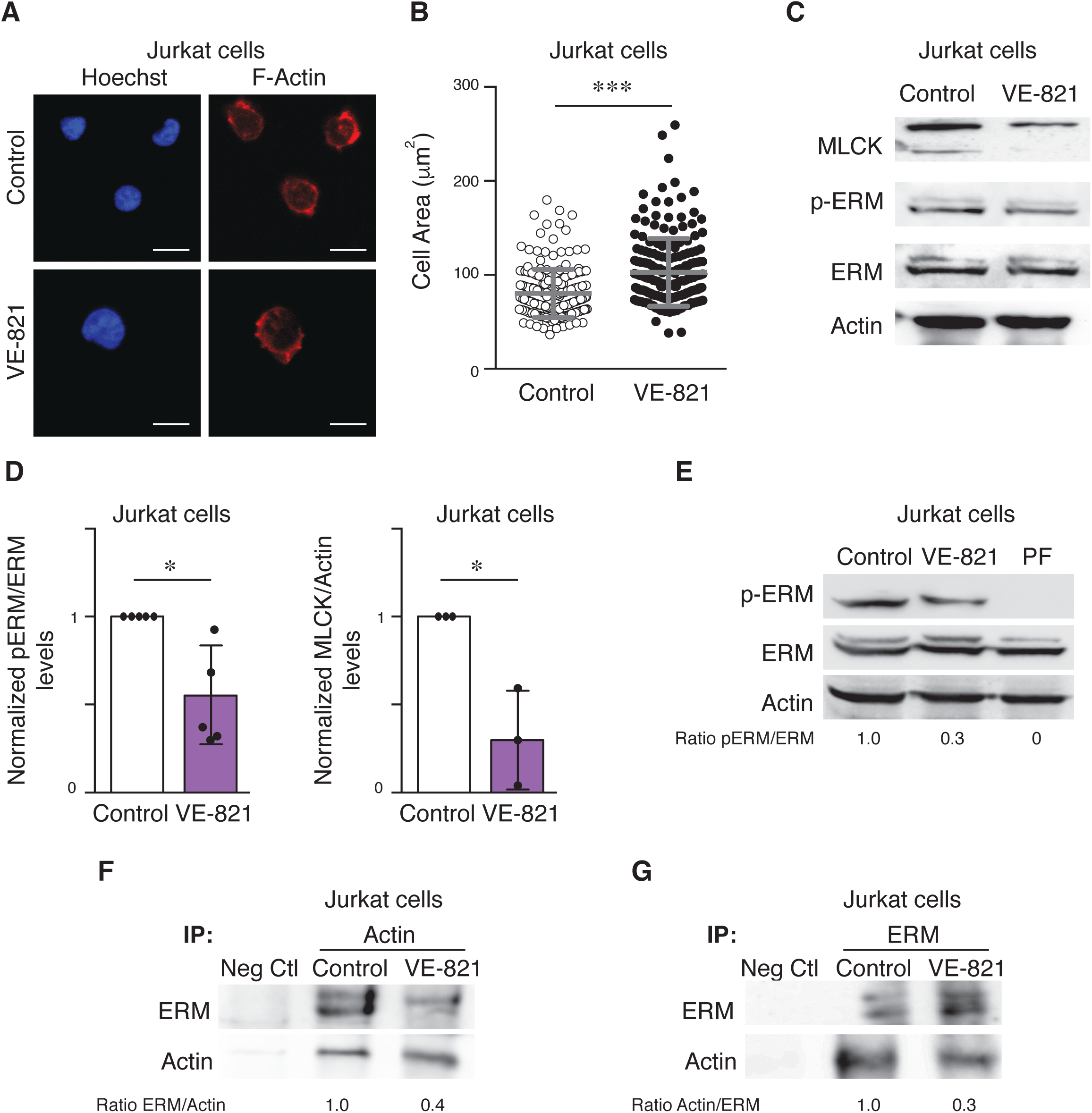
ATR inhibition alters the cortical actomyosin cytoskeleton of T-ALL. **(A)** Representative immunofluorescence images of control and VE-821-treated Jurkat cells cultured on poly-L-lysine-coated slides for 1 h, fixed, and stained with phalloidin (red) and Hoechst (blue) (scale bar: 10 µm). **(B)** Cellular area quantification from phalloidin-staining (*n* = 210-193 cells; *N* = 3). *** P<0.001. **(C)** Representative immunoblots for MLCK and phospho-ERM proteins in whole-cell lysates of control and VE-821-treated Jurkat cells. **(D)** phospho-ERM/ERM and MLCK/Actin ratios. (*N* = 3-5). **(E)** Representative immunoblots showing phospho-ERM levels in total lysates from control and Chk-1-inhibited Jurkat cells. **(F)** Representative immunoblots showing actin–ERM interactions following immunoprecipitation of ERM from control and VE-821-treated Jurkat cell lysates. Actin was detected by immunoblotting in ERM immunoprecipitation. Neg. Ctl. represents the control IP, and numbers represent the protein quantification ratio after normalizing control cell values to 1. **(G)** Representative immunoblots showing actin–ERM interactions following immunoprecipitation of actin from control and VE-821-treated Jurkat cell lysates. ERM was detected from actin immunoprecipitation as in (F).

## DISCUSSION

Here, we described that ATR inhibition has opposite effects on migration through collagen and across endothelial monolayers in T-ALL cells. The extracellular environment determines the DNA damage response of tumor cells (35), including leukemic cells that have to adapt the nucleus and the cytoskeleton to different mechanical environments (36,37). The increased nuclear deformation and changes in MLCK expression, ERM phosphorylation and ERM–actin association observed after ATR inhibition suggest that these migratory outcomes involve different nuclear and cortical requirements. Thus, our results indicate that ATR might coordinate different mechanical requirements during migration, suggesting that a greater capacity for nuclear deformation does not necessarily improve migration across all physical barriers.

Nuclear mechanical properties are determined by the combined contribution of the nuclear lamina, chromatin organization and the coupling between the nucleus and the cytoskeleton (38,39). DNA damage response is critical in human diseases (40,41), and the enzymes regulating this response have been proposed as targets for cancer therapy, including leukemia (42–45). While these works indicate the effect of ATR targeting as genotoxic agent alone or in combination with other chemotherapies, our results show that genetic or pharmacological inhibition of ATR increased nuclear area and altered nuclear shape in both T-ALL cell lines and in primary T-ALL cells, together with changes in chromatin accessibility and nuclear mechanical behaviour. Importantly, we analyzed nuclear morphology and migration at time points and concentrations at which cell cycle distribution and viability were not substantially altered. The similar nuclear phenotype observed after pharmacological inhibition and genetic depletion supports a contribution of ATR to these changes.

Interactions between DNA damage, chromatin changes and nuclear stiffness have been extensively reported (46–48). Although ATR-targeted cells displayed increased nuclear area, ATR inhibition increased the apparent indentation modulus and surface heterogeneity of isolated nuclei while also increasing their deformation under mechanical compression. These assays apply different loading conditions and do not support a simple description of the nucleus as uniformly softer or stiffer. AFM provides a local estimate of the resistance of isolated nuclei to indentation, whereas mechanical compression measures changes in nuclear shape upon spatial restriction. This interpretation is consistent with studies showing that AFM might reflect different mechanical behaviours (49), and that acute chromatin decompaction can affect nuclear morphology and mechanical behaviour by increasing the nuclear stiffness (50). Although previous studies have shown that ATR contributes to the cellular response to mechanical stress and nuclear adaptation (14,23,51), our results indicate that ATR alters the nuclei of T-ALL cells, increasing their deformation under confinement despite an increased apparent indentation modulus in isolated nuclei.

In contrast to the increased heterochromatin reported in ATR-defective U2OS and HeLa cells (14), ATR inhibition in leukemic cells increased H3K4me3 levels and DNase I sensitivity. Furthermore, although previous literature suggests that activation of H3K4 methylation might lead to an inactivation of ATR (52,53), we found that ATR inhibition increased chromatin accessibility, WDR5 association with chromatin and global H3K4me3 levels in ALL cells. Previous work implicated WDR5-dependent H3K4 methylation in nuclear adaptation to 3D environments and cell motility (22,32), making this epigenetic change a plausible link between ATR inhibition and the nuclear changes observed. While it remains to be determined whether these epigenetic alterations impact additional cellular functions, our results suggest that ATR targeting might be connected to H3K4 tri-methylation and WDR5 location at the chromatin.

The nuclear changes observed were accompanied by increased DNase I sensitivity, suggesting that chromatin became more accessible following ATR inhibition. Given that ATR regulates gene transcription in response to mechanical stimuli to modulate cell differentiation (54,55), it is remarkable that ATR inhibition resulted in a predominance of downregulated genes in both cell lines, despite the observed increases in global H3K4me3 levels and DNase I sensitivity. ATR has been implicated in the regulation of RNA polymerase II during S phase (56,57), and H3K4me3 can regulate promoter-proximal pause-release and elongation (58). These findings provide plausible explanations for the relationship between chromatin and mRNA changes observed. It has been reported that mechanical stress drives transcriptional changes and cellular reprogramming (59,60), and our comparison between Jurkat and CCRF-CEM cells showed shared representation of mitochondrial and ribosomal categories in both cell lines, together with cell-line-specific changes, indicating that the consequences of ATR inhibition are influenced by cellular context. These findings rise the possibility that targeting ATR might affect mitochondrial/ribosomal component programs required for T-ALL cells to tolerate and adapt to mechanical and replicative stress. This observation is particularly relevant considering the metabolic requirements of proliferating T-ALL cells and the relationship between ATR signalling, nucleotide availability and replication stress (19). Similarly, the enrichment of ribosomal components may reflect changes in protein synthesis-associated processes during the response to treatment. Whether these changes translate into altered migratory capacity of T-ALL cells in confined conditions remains to be established.

Of note, library-size-normalized ChIP-seq profiles did not reveal widespread increases in H3K4me3 signal intensity. This apparent discrepancy highlights that while immunoblotting determined the global H3K4me3 levels, normalized ChIP-seq measured relative enrichment of H3K4me3 across genomic regions. H3K4 methylation is a typical marker for transcriptional activity (61); although additional functions independent of gene expression have been described (62,63). Changes in mRNA might involve transcription initiation, elongation or RNA stability, but our integrated multi-omic profiles showed that changes in H3K4me3 density are not accompanied by a linear correspondence in steady-state mRNA abundance. This suggests that the gene expression response to ATR inhibition cannot be explained solely by the localized H3K4me3 dynamics.

ATR interference increased migration through confined 3D matrices, consistent with the idea that increased nuclear deformability facilitates movement through narrow spaces (64). An important finding of this study is that ATR inhibition increased migration in confined 3D collagen while reducing the number of leukemic cells recovered from peripheral tissues 3 h after intravenous injection. This assay indicates an early distribution rather than long-term colonization; therefore, we cannot discard that ATR inhibition also affects later stages of leukemic dissemination. During the metastatic process tumor cells have to navigate across multiple physical barriers (65), and the reduction in transendothelial migration of ATR-inhibited cells provides a plausible cellular explanation for the lower recovery of leukemic cells. It is possible that a more deformable nucleus may not be sufficient to overcome the requirements to interact with the vascular endothelium and generate the forces required for an effective diapedesis (66).

Transcriptomic analysis identified changes in cytoskeletal and adhesion-associated genes following VE-821 treatment may help explain why increased collagen invasion was accompanied by reduced transendothelial migration. The cytoskeleton is critical to promote chromatin and nuclear changes induced by biophysical stimuli (67–69), however, we detected no substantial changes in the expression of integrins, chemokine receptors, total F-actin levels or static adhesion to VCAM-1 of T-ALL cells upon VE-821 treatment. ERM proteins participate in the coupling between the plasma membrane and cortical actin, whereas MLCK contributes to actomyosin contractility (70,71). The reduction in MLCK expression, ERM phosphorylation and ERM–actin association therefore suggests altered cortical regulation rather than a general loss of actin or adhesion receptors. Pharmacological MLCK inhibition with ML-7 reproduced the increased spreading observed after ATR inhibition, suggesting the involvement of MLCK on cell spreading. The increased spreading observed upon 2D substrates also suggests that ATR inhibition does not simply suppress all aspects of cell movement or adhesion.

MLCK activity and ERM-actin interactions have been extensively linked to active transendothelial migration of leukocytes (72); making them plausible contributors to the phenotype observed. Furthermore, ERM proteins have distinct functions during 2D and 3D migration (73), which may contribute to the differences observed. The effect of ATR inhibition may therefore depend on the balance between nuclear deformation and cortical functions in each environment. Under confined 3D conditions, the increased nuclear deformability may facilitate passage through restrictive spaces. During transendothelial migration, however, efficient movement requires the coordination of nuclear deformation with membrane–cortex interaction, adhesion and contractile forces (74,75). A defect in these processes could therefore limit vascular transmigration despite the increased deformability of the nucleus. Comparison with Chk-1 inhibition further suggests that the nuclear and cortical responses differ in their dependence on canonical ATR–Chk-1 signalling. Although ATR and Chk-1 act in the same major replication stress response pathway, they can also have distinct functions (34,76). Under the conditions tested, Chk-1 inhibition did not reproduce the nuclear deformation or H3K4 tri-methylation responses to ATR inhibition, although both treatments reduced ERM phosphorylation. These findings suggest that some cortical effects may overlap, whereas the nuclear response may involve additional ATR-dependent components. Further work will be required to determine which ATR-dependent signalling components connect replication-associated stress with chromatin remodeling and cytoskeletal regulation in T-ALL cells.

In summary, our findings identify ATR as a regulator of nuclear and cytoskeletal adaptation in T-ALL cells, affecting their migration in a context-dependent manner. ATR inhibition alters nuclear architecture and mechanical behaviour and promotes migration through confined 3D collagen matrices, whereas impairing transendothelial migration and reducing early leukemic cell recovery from peripheral tissues. These responses are accompanied by changes in chromatin accessibility, and H3K4me3 levels, while changes in MLCK and ERM regulation suggest altered cortical response. Together, these findings show that increased nuclear deformability alone is not sufficient to ensure efficient leukemic dissemination and highlight the distinct mechanical requirements of migration through extracellular matrices and across the vascular endothelium.

## DATA AVAILABILITY

All sequencing data from this study will be deposited in the National Center for Biotechnology Information Gene Expression Omnibus (GEO) and will be accessible through the GEO Series under accession number. All other relevant data are available within the article and supplementary files, or from the corresponding author upon request.

## ACKNOWLEDGEMENTS

The authors thank the Microscopy Unit of Instituto de Investigación Biosanitaria Gregorio Marañón (IiSGM) and the Microscopy Facility of CIB Margarita Salas for assistance with confocal analyses, and to the Bioinformatics and Animal Facilities of the CIB Margarita Salas for their assistance and support with the RNAseq and ChIPseq analyses and the in vivo experiments. We also thank Patricia Pedraz and Cristina Flors for assistance and support at AFM Facility in IMDEA Nanomaterials. Finally, we thank the support of the CSIC-Hub Conexión Cáncer.

## AUTHORS INFORMATION

### CONTRIBUTIONS

M.A. was involved in experimental design, conducted experiments and contributed to data interpretation and discussion. N.B., Y.S., and G.P., performed experiments and data analysis. E.A.L., and I.B., supervised the experiments related to transendothelial migration and contributed to data interpretation. E.G.S., A.B., and M.R., contributed to clinical analyses and primary sample experiments. J.R.M. was involved in study conception and experimental design, data analysis, and writing the manuscript with editorial assistance from the other authors.

## ETHICS DECLARATIONS

### ETHICAL APPROVAL

This study was reviewed and approved by the CSIC and Hospital Niño Jesús ethics committee, and written informed consent was obtained from all parents, according to the Helsinki Declaration of 1975. All experiments involving animals were approved by the OEBA (Organ for Evaluating Animal Wellbeing) at CIB Margarita Salas and Madrid Regional Department of Environment, with reference PROEX 228.4/21. All procedures for animal experiments were approved by the Committee on the Use and Care of Animals and carried out in strict accordance with the institution guidelines and the European and Spanish legislations for laboratory animal care.

### CONSENT FOR PUBLICATION

All authors have read the manuscript and agreed to its publication.

## REFERENCES

[1] A. J. Lomakin, et al., Science 2020, 370, eaba2894. 10.1126/science.aba2894

[2] V. Venturini, et al., Science 2020, 370, eaba2644. 10.1126/science.aba2644

[3] P. Friedl, S. Alexander, Cell 2011, 147, 992. 10.1016/j.cell.2011.11.016

[4] H. Zhou, et al., Cancer Commun. 2022, 42, 374. 10.1002/cac2.12294

[5] K. Wolf, et al., J. Cell Biol. 2013, 201, 1069. 10.1083/jcb.201210152

[6] K. Jiang, S. B. Lim, J. Xiao, D. S. Jokhun, M. Shang, X. Song, P. Zhang, L. Liang, B. C. Low, G. V. Shivashankar, C. T. Lim, Adv. Sci. 2023, 10, 2201663. 10.1002/advs.202201663

[7] Y. Kalukula, et al., Nat. Rev. Mol. Cell Biol. 2022, 23, 583. 10.1038/s41580-022-00480-z

[8] G. P. de Freitas Nader, A. Williart, M. Piel, Curr. Opin. Cell Biol. 2021, 72, 137. 10.1016/j.ceb.2021.07.008

[9] M. Raab et al., Science 2016, 352, 359. 10.1126/science.aad7611

[10] C. M. Denais et al., Science 2016, 352, 353. 10.1126/science.aad7297

[11] E. J. Brown, D. Baltimore, Genes Dev. 2003, 17, 615. 10.1101/gad.1067403

[12] F. Esposito, S. Francia, J. Cell Sci. 2026, 139, jcs263423. 10.1242/jcs.263423

[13] I. Chiolo, et al., Nat. Rev. Mol. Cell Biol. 2025, 26, 538. 10.1038/s41580-025-00828-1

[14] G. R. Kidiyoor, et al., Nat. Commun. 2020, 11, 4828. 10.1038/s41467-020-18580-9

[15] G. R. Kidiyoor, A. Kumar, M. Foiani, DNA Repair 2016, 44, 143. 10.1016/j.dnarep.2016.05.020

[16] S. E. Fordham, et al., Blood Adv. 2018, 2, 1157. 10.1182/bloodadvances.2017015214

[17] R. Jossé, et al., Cancer Res. 2014, 74, 6968. 10.1158/0008-5472.CAN-13-3369

[18] R. Prevo, et al., Cancer Biol. Ther. 2012, 13, 1072. 10.4161/cbt.21093

[19] T. M. Le, et al., Nat. Commun. 2017, 8, 241. 10.1038/s41467-017-00221-3

[20] E. A. Raetz, D. T. Teachey, Hematology Am. Soc. Hematol. Educ. Program 2016, 2016, 580. 10.1182/asheducation-2016.1.580

[21] E. Madrazo et al., Oncogene 2022, 41, 1324. 10.1038/s41388-021-02168-8

[22] R. González-Novo, et al., Eur. J. Cell Biol. 2023, 102, 151343. 10.1016/j.ejcb.2023.151343

[23] A. Kumar et al., Cell 2014, 158, 633. 10.1016/j.cell.2014.05.046

[24] H. Su, et al., Transl. Oncol. 2023, 36, 101743. 10.1016/j.tranon.2023.101743

[25] A. D. Stephens, et al., Mol. Biol. Cell 2018, 29, 220. 10.1091/mbc.E17-06-0410

[26] R. González-Novo, et al., Cell Rep. Phys. Sci. 2026, 7, 103116. 10.1016/j.xcrp.2026.103116

[27] A. L. McGregor, C.-R. Hsia, J. Lammerding, Curr. Opin. Cell Biol. 2016, 40, 32. 10.1016/j.ceb.2016.01.011

[28] N. Reymond, B. B. d’Água, A. J. Ridley, Nat. Rev. Cancer 2013, 13, 858. 10.1038/nrc3628

[29] A. J. Maniotis, C. S. Chen, D. E. Ingber, Proc. Natl. Acad. Sci. U. S. A. 1997, 94, 849. 10.1073/pnas.94.3.849

[30] J.-H. Lee, et al., J. Cell Biol. 2004, 167, 327. 10.1083/jcb.200403091

[31] G. Totsukawa, et al., J. Cell Biol. 2004, 164, 427. 10.1083/jcb.200306172

[32] P. Wang, et al., Proc. Natl. Acad. Sci. U. S. A. 2018, 115, 8581. 10.1073/pnas.1719405115

[33] Q. Liu, et al., Genes Dev. 2000, 14, 1448.

[34] R. Buisson, J. L. Boisvert, C. H. Benes, L. Zou, Mol. Cell 2015, 59, 1011. 10.1016/j.molcel.2015.07.029

[35] M. Deng, et al., Sci. Adv. 2020, 6, eabb2630. 10.1126/sciadv.abb2630

[36] A. E. Whiteley, T. T. Price, G. Cantelli, D. A. Sipkins, Nat. Rev. Cancer 2021, 21, 461. 10.1038/s41568-021-00355-z

[37] A. Lyu, S. H. Nam, R. S. Humphrey, T. M. Horton, L. I. R. Ehrlich, Exp. Mol. Med. 2024, 56, 2337. 10.1038/s12276-024-01335-7

[38] A. Athirasala, N. Hirsch, A. Buxboim, Curr. Opin. Cell Biol. 2017, 46, 119. 10.1016/j.ceb.2017.04.004

[39] D. M. Graham, K. Burridge, Curr. Opin. Cell Biol. 2016, 40, 98. 10.1016/j.ceb.2016.03.006

[40] A. Marechal, L. Zou, Cold Spring Harb. Perspect. Biol. 2013, 5, a012716. 10.1101/cshperspect.a012716

41. S. P. Jackson, J. Bartek, Nature 2009, 461, 1071. 10.1038/nature08467

[42] L. M. Karnitz, L. Zou, Clin. Cancer Res. 2015, 21, 4780. 10.1158/1078-0432.CCR-15-0479

[43] S. De Mel, et al., Front. Oncol. 2024, 14, 1307839. 10.3389/fonc.2024.1307839

[44] J. Szydzik, et al., Nat. Commun. 2021, 12, 6813. 10.1038/s41467-021-27057-2

[45] M. Boudny, M. Trbusek, Cancer Treat. Rev. 2020, 88, 102026. 10.1016/j.ctrv.2020.102026

[46] M. M. Nava et al., Cell 2020, 181, 800. 10.1016/j.cell.2020.03.052

[47] C.-R. Hsia et al., iScience 2022, 25, 104978. 10.1016/j.isci.2022.104978

[48] Á. dos Santos, et al., Nucleic Acids Res. 2021, 49, 340. 10.1093/nar/gkaa1202

[49] C. M. Hobson, et al., Mol. Biol. Cell 2020, 31, 1788. 10.1091/mbc.E20-01-0073

[50] A. Mitra, et al., Proc. Natl. Acad. Sci. U. S. A. 2025, 122, e2416659122. 10.1073/pnas.2416659122

[51] M. Chatzifrangkeskou, et al., Sci. Adv. 2025, 11, eadr5683. 10.1126/sciadv.adr5683

[52] B. K. Neilsen et al., BMC Cancer 2018, 18, 673. 10.1186/s12885-018-4580-6

[53] S. Y. Chong, et al., Nat. Commun. 2020, 11, 809. 10.1038/s41467-020-14595-4

[54] T. Roy, et al., Nucleic Acids Res. 2025, 53, gkaf852. 10.1093/nar/gkaf852

[55] X. Tu, et al., J. Clin. Invest. 2026, 136, e192285. 10.1172/JCI192285

[56] J. Wang, S. Pathak, M. Jones, J. Mao, M. Saponaro, iScience 2026, 29, 116663. 10.1016/j.isci.2026.116663

[57] H. B. Landsverk, L. E. Sandquist, S. C. Sridhara, G. E. Rødland, J. C. Sabino, S. F. de Almeida, B. Grallert, L. Trinkle-Mulcahy, R. G. Syljuåsen, Nucleic Acids Res. 2019, 47, 1797. 10.1093/nar/gky1233

[58] H. Wang, Z. Fan, P. V. Shliaha, M. Miele, R. C. Hendrickson, X. Jiang, K. Helin, Nature 2023, 615, 339. 10.1038/s41586-023-05780-8

[59] X. Gao, Y. Li, J. W. N. Lee, J. Zhou, V. Rangaraj, J. Marlena, A. W. Holle, Adv. Sci. 2025, 12, 2415407. 10.1002/advs.202415407

[60] S. M. Park, J. H. Lee, K. S. Ahn, H. W. Shim, J. Y. Yoon, J. Hyun, J. H. Lee, S. Jang, K. H. Yoo, Y. K. Jang, T. J. Kim, H. K. Kim, M. R. Lee, J. H. Jang, H. Shim, H. W. Kim, Adv. Sci. 2023, 10, 2303395. 10.1002/advs.202303395

[61] H. Santos-Rosa et al., Nature 2002, 419, 407. 10.1038/nature01080

[62] F. S. Howe, H. Fischl, S. C. Murray, J. Mellor, BioEssays 2017, 39, 1. 10.1002/bies.201600095

[63] H. Wang, K. Helin, Trends Cell Biol. 2025, 35, 115. 10.1016/j.tcb.2024.06.001

[64] X. Liu, et al., Adv. Sci. 2026, 13, e75470. 10.1002/advs.75470

[65] A. W. Lambert, D. R. Pattabiraman, R. A. Weinberg, Cell 2017, 168, 670. 10.1016/j.cell.2016.11.037

[66] A. Dupas, J. G. Goetz, N. Osmani, J. Cell Sci. 2024, 137, jcs262066. 10.1242/jcs.262066

[67] H.-R. Thiam, et al., Nat. Commun. 2016, 7, 10997. 10.1038/ncomms10997

[68] E. M. Hatch, M. W. Hetzer, J. Cell Biol. 2016, 215, 27. 10.1083/jcb.201603053

[69] J. Zhang et al., Small 2020, 16, 1907688. 10.1002/smll.201907688

[70] G. Salbreux, G. Charras, E. Paluch, Trends Cell Biol. 2012, 22, 536. 10.1016/j.tcb.2012.07.001

[71] R. G. Fehon, A. I. McClatchey, A. Bretscher, Nat. Rev. Mol. Cell Biol. 2010, 11, 276. 10.1038/nrm2866

[72] W. A. Muller, Annu. Rev. Pathol. 2011, 6, 323. 10.1146/annurev-pathol-011110-130224

[73] A. Estecha, et al., J. Cell Sci. 2009, 122, 3492. 10.1242/jcs.053157

[74] X. Cao, et al., Biophys. J. 2016, 111, 1541. 10.1016/j.bpj.2016.08.011

[75] A. C. I. van Steen, W. J. van der Meer, I. E. Hoefer, J. D. van Buul, Atherosclerosis 2020, 315, 102. 10.1016/j.atherosclerosis.2020.06.004

[76] J. Speroni, M. B. Federico, S. F. Mansilla, G. Soria, V. Gottifredi, Proc. Natl. Acad. Sci. U. S. A. 2012, 109, 7344. 10.1073/pnas.1116345109

