## Supplemental Info for "ATR coordinates nuclear and cytoskeletal mechanics during leukemic migration"

### SUPPLEMENTARY MATERIALS AND METHODS

| Antibodies | Host | Company | Cat. No. |
|---|---|---|---|
| anti-ATR | Rabbit | Cell Signaling | 2790S |
| anti-Actin | Rabbit | AbCam | EPR16769 |
| anti-ASH2L | Rabbit | Cell Signaling | 5019S |
| anti-CXCR4 (4G10) | Mouse | Santa Cruz Biotech | K0325 |
| anti-ERM | Rabbit | EMD Millipore Corp. | AB3843 |
| anti-H3 | Rabbit | Cell Signaling | 4499L |
| anti-H3K4me3 | Rabbit | EMD Millipore Corp. | 05-745R |
| anti-Integrin-α4 (HP2/1) | Mouse | Santa Cruz Biotech | sc-365209 |
| anti-Integrin-ß1 (12G10) | Mouse | Santa Cruz Biotech | sc-59827 |
| anti-Integrin-ß1 (TS2/16) | Mouse | Santa Cruz Biotech | sc-53711 |
| anti-Lamin B1 (C-5) | Mouse | Santa Cruz Biotech | sc-365962 |
| anti-MLCK | Rabbit | Sigma-Aldrich | SAB1300116 |
| anti-Mouse Secondary HRP | Goat | Bio-Rad | 1706516 |
| anti-Mouse-Alexa Fluor 488 | Goat | Invitrogen | A11001 |
| anti-phospho-ERM | Rabbit | Sigma-Aldrich | SAB4504260 |
| anti-phospho-ERM | Rabbit | Merck | AB3832 |
| anti-Rabbit Secondary HRP | Goat | Fisher Scientific | 15217664 |
| anti-RhoGDI-α (A-20) | Mouse | Santa Cruz Biotech | sc-373724 |
| anti-WDR5 (G-9) | Mouse | Santa Cruz Biotech | sc-393080 |

| Materials and reagents | Company | Cat. No. |
|---|---|---|
| Agarose | PanReac | A8963 |
| AmershaM Protran 0.45 NC membranes | GE Healthcare Life Sci | 10600002 |
| ATR shRNA Plasmid | Santa Cruz Biotech | sc-29763-SH |
| Bovine Serum Albumin (BSA) | PanReac | A6588 |

| | | |
|---|---|---|
| CellAdhere Type I Collagen, Bovine | StemCell Technologies | 7001 |
| CellTrace CFSE | Thermo Fisher Sci | C34554 |
| CellTrace Far Red | Thermo Fisher Sci | C34564A |
| CellTrace Violet | Thermo Fisher Sci | C34557 |
| CellTrace Yellow | Thermo Fisher Sci | C34573 |
| CXCL12/SDF - 1α | PeproTech | 300-28A |
| Control shRNA Plasmid A | Santa Cruz Biotech | sc-108060 |
| DABCO mounting medium | Sigma-Aldrich | 10981 |
| Dimethyl Sulfoxide - DMSO | Calbiochem | 317275 |
| Dithiothreitol (DTT) | Bio-Rad | 1610610 |
| DNase I | Thermo Fisher Sci | EN0521 |
| ECL Western Blotting Substrate | Westar Sun | XLS063L |
| EGM SingleQuots | Lonza | CC-4133 |
| Endothelial Cell Growth Basal Medium (EBM) | Lonza | CC-3121 |
| Fetal Bovine Serum (FBS) | Sigma-Aldrich | 12103C |
| FITC Annexin V Apoptosis Detection Kit with PI | ImmunoStep | ANXVKF-100T |
| Formaldehyde solution min 35% | Merck | 1040001 |
| Glass bottom dishes - Low (35 mm) | Ibidi | 80136 |
| Glass bottom dishes (50x7 mm) | Willco | GWST-5040 |
| Hoechst 33342 | Invitrogen | H3570 |
| Human VCAM-1 | PeproTech | 150-04 |
| HUVEC | Lonza | K3CC-2519 |
| Laemmli SDS sample buffer (6X) | Thermo Fisher Sci | 15493939 |
| ML-7 | Sigma-Aldrich | ST9H93D8DBDB |
| Nonidet P-40 (NP-40) | PanReac | A1694 |
| OICR-9429 | Tocris Bioscience | 5267 |
| Phalloidin-Alexa Fluor 568 | Invitrogen | A12380 |

| | | |
|---|---|---|
| Phalloidin-CF488A | Biotium | 00042 |
| Paraformaldehyde | Merck | 4005 |
| PF-477736 | Sigma-Aldrich | PZ0186 |
| Poly-L-Lysine | Sigma-Aldrich | P1399 |
| Propidium Iodide | Sigma-Aldrich | P4864 |
| Protease Inhibitor Cocktail | Sigma-Aldrich | 535142-1ML |
| Proteinase K | Sigma-Aldrich | P1308 |
| Protein A/G PLUS-Agarose | Santa Cruz Biotech | sc-2003 |
| RNase A | Sigma-Aldrich | R4875 |
| RPMI 1640 Medium | Gibco | 11530586 |
| SimpleChIP Plus Sonication ChIP Kit | Cell Signaling | 56383 |
| SYBR Green I | Sigma-Aldrich | S9430 |
| Corning Transwell multiple well plate | Corning | 10107341 |
| TRI Reagent® | Sigma-Aldrich | T9424 |
| Triton X-100 | Fisher Scientific | 150232OB-100 |
| UltraCruz® Transfection Reagent | Santa Cruz Biotech | sc-395739 |
| VE-821 | Sigma-Aldrich | SML1415 |
| β-mercaptoethanol | Sigma-Aldrich | M3148 |

**Immunoprecipitation.** $7 \times 10^6$ cells under different conditions were lysed in actin-binding buffer (50 mM Tris-HCl [pH 7.4], 150 mM NaCl, 0.2% NP-40, 2 mM $MgCl_2$) supplemented with protease inhibitor cocktail. Lysates were vortexed, incubated on ice for 10 min, and centrifuged at 13,600 x g for 5 min at 4 °C. After preclearing with Protein A/G agarose beads, supernatants were incubated with antibodies (3 h, 4 °C), followed by coupling to Protein A/G agarose beads (overnight, 4 °C) on rotation. After washing, protein complexes were eluted with Laemmli Buffer and boiled at 95 °C for 5 min. Eluted proteins were analyzed by immunoblotting as described above.

**DNAse I assay.** 5 × $10^6$ isolated nuclei per condition were resuspended in DNAse buffer (Tris-HCl 10 mM [pH 7.5], $MgCl_2$ 2.5 mM, $CaCl_2$ 0.5 mM) and digested with 0.5 U of DNAse I (15 min, 25 °C). Reactions were stopped by adding STOP buffer (10 μM Tris-HCl, 5 mM EDTA, 200 mM NaCl, 0.2% SDS), and samples were subsequently incubated with 1 μg RNAse A (30 min, 37 °C) and 1 μg Proteinase K (overnight at 52 °C). Digested DNA was isolated with phenol/chloroform and precipitated with 2.5 volumes of ethanol and 0.1 volume of 3 M sodium acetate at −20°C overnight. The DNA pellet was dissolved in water, resolved on a 1.5% (w/v) agarose gel containing SYBR green and visualized in a ChemiDoc. DNA digestion profiles were quantified using ImageJ.

**Chromatin Immunoprecipitation-Sequencing (ChIP-seq).** 7 × $10^7$ cells under different conditions were harvested, and Chromatin Immunoprecipitation (ChIP) was performed following the manufacturer's protocol of the SimpleChIP® Plus Sonication Chromatin IP Kit. Cells were crosslinked with formaldehyde, lysed, and chromatin was fragmented by sonication into 100–300 bp fragments. Subsequently, chromatin was immunoprecipitated using 3 μL of rabbit anti-H3K4me3 and the Protein G magnetic beads provided in the kit. Chromatin was then eluted, reverse crosslinking was performed, and DNA was purified. Two independent biological replicates per condition were analyzed in Jurkat and CCRF-CEM cells, and no input DNA controls were used. Purified ChIP DNA was shipped to BGI Genomics for library construction (end-repair, adapter ligation, PCR amplification, and single-stranded circularization), and high-throughput sequencing on the DNBSEQ platform (50-bp single-end reads). Downstream data processing and bioinformatic analysis were performed at the CIB-CSIC Bioinformatics and Biostatistics Unit. Raw reads were filtered using SOAPnuke v2.2.6 (1). Reads were discarded if they contained adapter sequences, if more than 50% of their bases had Phred quality scores below 10, or if ambiguous nucleotides (N) constituted more than 1% of their bases. Filtered reads were aligned to the human reference genome, GRCh38 (hg38), using Bowtie2 v2.4.5 (2). Duplicate reads were excluded, and only alignments with mapping quality scores greater than 20 were retained. H3K4me3-enriched regions were identified using MACS3 v3.0.0b1 (3) with default parameters. Peaks were annotated to genomic features and associated genes using ChIPseeker v1.30.3 (4,5) and the GENCODE release 29 annotation for GRCh38 (6). Differential H3K4me3 enrichment between VE-821-treated and control samples was assessed separately for each cell line using PePr v1.1.24 (7) in differential mode with default parameters, including both biological replicates per condition. PePr models read counts using a local negative binomial framework to account for variability among replicates. Differentially enriched regions were annotated using ChIPseeker and GENCODE release 29 to identify associated genes for downstream analyses. Genes associated with differentially enriched H3K4me3 regions were subjected to Gene Ontology (GO) (8,9) and Kyoto Encyclopedia of Genes and Genomes (KEGG) pathways (10,11,12) overrepresentation analyses using an upper-tailed hypergeometric test, with whole-genome genes serving as the background. Multiple-testing correction was performed using the Benjamini–Hochberg procedure, applied separately across all tested GO terms and all tested KEGG pathways. Terms and pathways with a false discovery rate (FDR) $< 0.05$ were considered significantly enriched. High-resolution candidate sites were identified using SeqSite v1.1.2 (13), and the corresponding genomic sequences were analyzed using MEME-ChIP v5.5.9 (14) against the human HOCOMOCOv11_core motif database (15) to detect enriched motifs potentially associated with transcription factor binding. ChIP-seq files used in this study will be deposited at the NCBI Gene Expression Omnibus (GEO).

**RNA Sequencing (RNA-seq).** Total RNA was extracted from $5 \times 10^6$ cells using TRI reagent (30 min, RT), followed by chloroform extraction (30 min, RT), isopropanol precipitation (15 min, 4 °C) and ethanol washing. Total RNA was resuspended in RNAsefree water, and purity and concentration were determined by NanoDrop ND-1000 Spectrophotometer. RNA samples from three independent biological replicates per condition (from Jurkat and CCRF-CEM cells) were shipped to BGI for library construction and sequencing. Libraries were generated by BGI (poly(A) mRNA enrichment, double-stranded cDNA synthesis, and single-stranded circular DNA library preparation) and sequenced on the BGI DNBSEQ platform (paired-end, 150 bp). Raw reads were filtered using SOAPnuke v1.5.6 (1), (parameters: -n 0.01 -l 15 -q 0.4) to remove reads containing adapter sequences, unknown bases (14) or low-quality bases. Filtered clean reads were aligned to the human reference genome (*Homo Sapiens* Ensembl release 105) using HISAT2 v2.2.1 (16) and to the reference gene set using Bowtie2 v2.5.0 (2). Gene expression was quantified using RSEM v1.3.1 software (17). Differentially expressed genes between control and VE-821-treated cells were identified in each cell line using DESeq2 v1.40.2 (18) (Q value ≤ 0.05). GO and KEGG enrichment analyses of differentially expressed genes were performed using the hypergeometric test (phyper function in 19) with a Q value ≤ 0.05. RNA-seq files used in this study will be deposited at the NCBI Gene Expression Omnibus (GEO).

**Cell adhesion.** 96-wells plates were coated with soluble recombinant human VCAM-1 at 2.5 μg/mL overnight at 4 °C. $3 \times 10^5$ cells were labelled with 5 μM BCECF-AM in RPMI (30 min). Then, cells were washed in 0.5% BSA in RPMI and seeded onto VCAM-1-coated wells for 30 min. Subsequently, cells were washed with 0.5 % BSA in RPMI, attached cells were lysed with 0.1 % SDS in PBS and fluorescence at 490 nm was quantified in a Varioskan (Thermo Fisher). Percentage of adherent cells from the total was calculated.

**Flow cytometry**. Cells were collected, washed with PBS (TBS for active β1 integrin) and kept on ice. Cells were sequentially incubated with primary antibodies (1 h, 4 °C), washed with PBS (or TBS), and incubated again with appropriate CF488A-conjugated secondary antibodies (1 h, 4 °C). Then, cells were resuspended in PBS, and analyzed on a CytoFLEX.

For each condition, a secondary antibody-only control and an unstained (autofluorescence) control were included. For intracellular staining of F-actin, cells were collected, washed with PBS, fixed, permeabilized, blocked and F-actin was stained with fluorochrome-conjugated phalloidin before their analysis on a CytoFLEX. Unstained cells were used as an autofluorescence control. The data obtained were analyzed using FlowJo software.

**Cell cycle.** Cells under different conditions were collected, washed in cold PBS and fixed in cold 70% (v/v) ethanol (−20 °C, 3 h). Then, cells were washed with PBS and incubated in PBS containing 0.1% Triton X-100, RNase A (100 µg/mL) and Propidium Iodide (50 µg/mL) (30 min, 37ºC). Then, cells were washed, resuspended in PBS, analyzed on CytoFLEX, and G1, S, and G2/M phases were determined.

**Apoptosis.** Cells under different conditions were collected, washed with PBS and incubated with the FITC Annexin V Apoptosis Detection Kit (ImmunoStep) and with Propidium Iodide (15 min, RT) before their analysis on a CytoFLEX. The proportion of live cells (Annexin V-FITC and Propidium Iodide negative) was calculated using FlowJo software.

**Transendothelial migration assay.** Transwell inserts (5 µm) were pre-coated with 0.1% gelatin in PBS. HUVECs were then seeded at a density of 10,000 cells per insert and grown for 4 days until a confluent monolayer was formed. To activate the endothelium, HUVEC monolayers were treated with 10 ng/mL TNF-α 18 h prior to the assay. Control and ATR-targeted cells were labelled with CellTrace Violet (1 µM) and CellTrace Yellow (2 µM), respectively (30 min); then, cells were mixed 1:1 and $3 \times 10^5$ cells were seeded onto HUVEC monolayers preactivated with 10 ng/mL TNF-α (18 h). RPMI supplemented with CXCL12 (150 ng/mL) was placed in the lower compartment. After 3 h, migrated cells were collected from the lower chamber, washed in PBS, and acquired on a CytoFLEX cytometer (Beckman Coulter). The percentage of cells stained with each vital label was quantified using FlowJo software.

**In-vivo short-term leukemia homing assay.** NOD-SCID-Il2rg-/- (NSG) mice (*Mus musculus*) were bred and maintained at the Servicio del Animalario del Centro de Investigaciones Biológicas Margarita Salas (CIB-CSIC) under number 28079-21A. All experiments involving animals are approved by the OEBA (Organ for Evaluating Animal Wellbeing) at CIB Margarita Salas and Madrid Regional Department of Environment, with reference PROEX 228.4/21. Control and ATR-inhibited cells were labelled with CellTracker Far Red (2 μM) and CFSE (5 μM), respectively (30 min); then, cells were mixed 1:1 and $5 \times 10^6$ cells of the mixture were injected intravenously in the tail vein of 9-week-old NSG mice. Dye-swap controls were not performed, as the labeling strategy had been extensively evaluated using reciprocal dye assignments in previous work (21,25). Animals were euthanized 3 h after injection, and the bone marrow from femurs, spleens, and livers were extracted, processed through mechanical disaggregation and filtered through 70 μm pores. Samples were incubated in erythrocyte lysis buffer (150 mM $NH_4Cl$, 10 mM $KHCO_3$, 0.1 mM EDTA) for 2 h at RT. Then, cells were washed in PBS and $1 \times 10^6$ events per tissue were acquired in a FACSCanto II cytometer. The percentage of labelled cells relative to the total number of events in the spleen, bone marrow and liver was analyzed using FlowJo software.

**Supplementary References**

1. Chen Y, Chen Y, Shi C, et al. SOAPnuke: a MapReduce acceleration-supported software for integrated quality control and preprocessing of high-throughput sequencing data. GigaScience. 2018;7(1):gix120. doi:10.1093/gigascience/gix120

2. Langmead B, Salzberg SL. Fast gapped-read alignment with Bowtie 2. Nat Methods. 2012;9(4):357–359. doi:10.1038/nmeth.1923

3. Doherty P, Hu Q, Zhuang Z, Zhang H, Penikalapati SC, Liu S, Liu T. MACS3: A Peak-calling Platform for Bulk and Single-cell Regulatory Genomics. Genomics Proteomics Bioinformatics. 2026;qzag097. doi:10.1093/gpbjnl/qzag097

4. Yu G, Wang LG, He QY. ChIPseeker: an R/Bioconductor package for ChIP peak annotation, comparison and visualization. Bioinformatics. 2015;31(14):2382–2383. doi:10.1093/bioinformatics/btv145

5. Wang Q, Li M, Wu T, et al. Exploring epigenomic datasets by ChIPseeker. Curr Protoc. 2022;2(10):e585. doi:10.1002/cpz1.585

6. Frankish A, Diekhans M, Ferreira AM, et al. GENCODE reference annotation for the human and mouse genomes. Nucleic Acids Res. 2019;47(D1):D766–D773. doi:10.1093/nar/gky955

7. Zhang Y, Lin YH, Johnson TD, et al. PePr: a peak-calling prioritization pipeline to identify consistent or differential peaks from replicated ChIP-Seq data. Bioinformatics. 2014;30(18):2568–2575. doi:10.1093/bioinformatics/btu372

8. Ashburner M, Ball CA, Blake JA, et al. Gene Ontology: tool for the unification of biology. Nat Genet. 2000;25(1):25–29. doi:10.1038/75556

9. The Gene Ontology Consortium. The Gene Ontology knowledgebase in 2026. Nucleic Acids Res. 2025;gkaf1292. doi:10.1093/nar/gkaf1292

10. Kanehisa M, Goto S. KEGG: Kyoto Encyclopedia of Genes and Genomes. Nucleic Acids Res. 2000;28(1):27–30. doi:10.1093/nar/28.1.27

11. Kanehisa M. Toward understanding the origin and evolution of cellular organisms. Protein Sci. 2019;28(11):1947–1951. doi:10.1002/pro.3715

12. Kanehisa M, Furumichi M, Sato Y, Matsuura Y, Ishiguro-Watanabe M. KEGG: biological systems database as a model of the real world. Nucleic Acids Res. 2025;53(D1):D672–D677. doi:10.1093/nar/gkae909

13. Wang X, Zhang X. Pinpointing transcription factor binding sites from ChIP-seq data with SeqSite. BMC Syst Biol. 2011;5(Suppl 2):S3. doi:10.1186/1752-0509-5-S2-S3

14. Machanick P, Bailey TL. MEME-ChIP: motif analysis of large DNA datasets. Bioinformatics. 2011;27(12):1696–1697. doi:10.1093/bioinformatics/btr189

15. Kulakovskiy IV, Vorontsov IE, Yevshin IS, et al. HOCOMOCO: towards a complete collection of transcription factor binding models for human and mouse via large-scale ChIP-Seq analysis. Nucleic Acids Res. 2018;46(D1):D252–D259. doi:10.1093/nar/gkx1106

16. Kim D, Langmead B, Salzberg SL. HISAT: a fast spliced aligner with low memory requirements. Nat Methods. 2015;12(4):357–360. doi:10.1038/nmeth.3317

17. Li B, Dewey CN. RSEM: accurate transcript quantification from RNA-Seq data with or without a reference genome. BMC Bioinformatics. 2011;12:323. doi:10.1186/1471-2105-12-323

18. Love MI, Huber W, Anders S. Moderated estimation of fold change and dispersion for RNA-seq data with DESeq2. Genome Biol. 2014;15(12):550. doi:10.1186/s13059-014-0550-8.

19. Kolde R. pheatmap: Pretty Heatmaps. R package version 1.0.12. 2019. https://CRAN.R-project.org/package=pheatmap

### SUPPLEMENTARY FIGURES

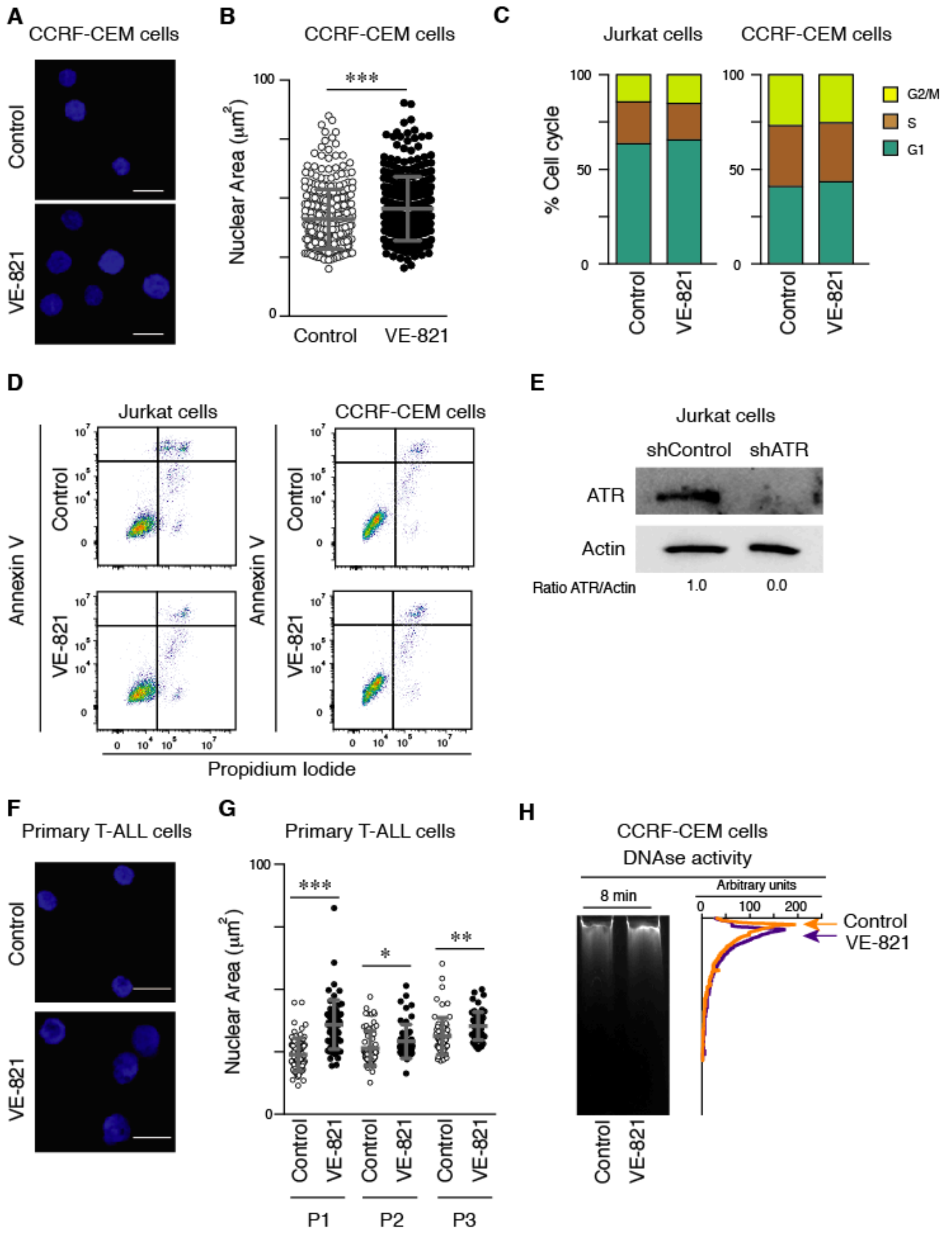


**Supplementary Figure S1**

**Supplementary Figure S1. ATR interference alters nuclear morphology without significantly affecting cell cycle progression or cell survival in different T-ALL cell models. (A)** CCRF-CEM cells were treated with or without VE-821 (1 µM for 24 h). Nuclear morphology was then visualized by Hoechst staining (DNA) (scale bar: 10 µm). **(B)** Nuclear area quantification (*n* = 292-293 cells; *N* = 3). *** P<0.001. **(C)** Cell cycle analysis of control and VE-821-treated Jurkat (left panel) and CCRF-CEM (right panel) cells stained with Propidium Iodide and analyzed by flow cytometry. **(D)** Apoptosis analysis of control and VE-821-treated Jurkat (left panels) and CCRF-CEM (right panels) cells by flow cytometry after Annexin V and Propidium Iodide staining. Representative dot plots showing the distribution of cells across four quadrants corresponding to viable cells, early apoptotic cells, late apoptotic cells, and necrotic cells. **(E)** Representative immunoblot showing ATR levels in total lysates from control (shControl) and ATR-depleted (shATR) Jurkat cells. **(F)** Nuclear morphology of control and VE-821-treated (10 μM for 24 h) primary T-ALL cells visualized by Hoechst staining (scale bar: 10 μm). **(G)** Nuclear area quantification (*n* = 52-79 nuclei). * P<0.05, ** P<0.01, *** P<0.001. **(H)** Representative gel image and corresponding relative-intensity profile of control and VE-821-treated CCRF-CEM cells after 15 min DNase I treatment. Arrowheads indicate the highest DNA-digestion peak.

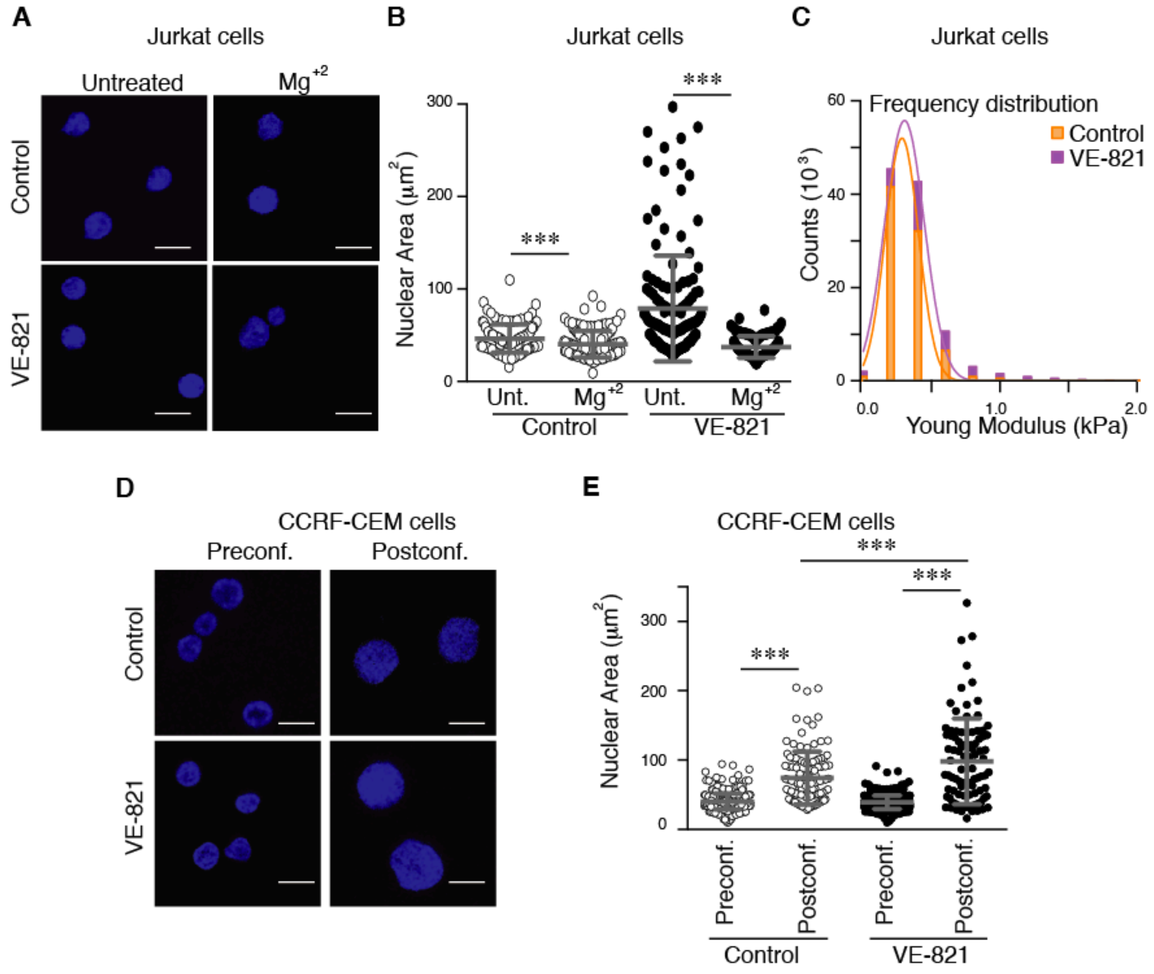


**Supplementary Figure S2**

**Supplementary Figure S2. ATR inhibition alters nuclear biomechanical properties and increases nuclear deformation under mechanical compression. (A)** Nuclear morphology of control and VE-821-treated Jurkat cells visualized by Hoechst staining in the presence or absence of $Mg^{2+}$ (scale bar: 10 µm). **(B)** Nuclear area quantification in the presence or absence of $Mg^{2+}$ ($n$ = 120-141 nuclei; $N$ = 2). **(C)** Frequency distribution of Young's modulus measurements obtained by atomic force microscopy (AFM) in control and VE-821-treated Jurkat cells. Counts are plotted against Young's modulus. **(D)** Representative images of control and VE-821-treated CCRF-CEM cells under mechanical compression visualized by Hoechst staining (scale bar: 10 µm). **(E)** Nuclear area quantification before and after mechanical compression ($n$ = 493/117-507/93 nuclei; $N$ = 2). *** P<0.001.

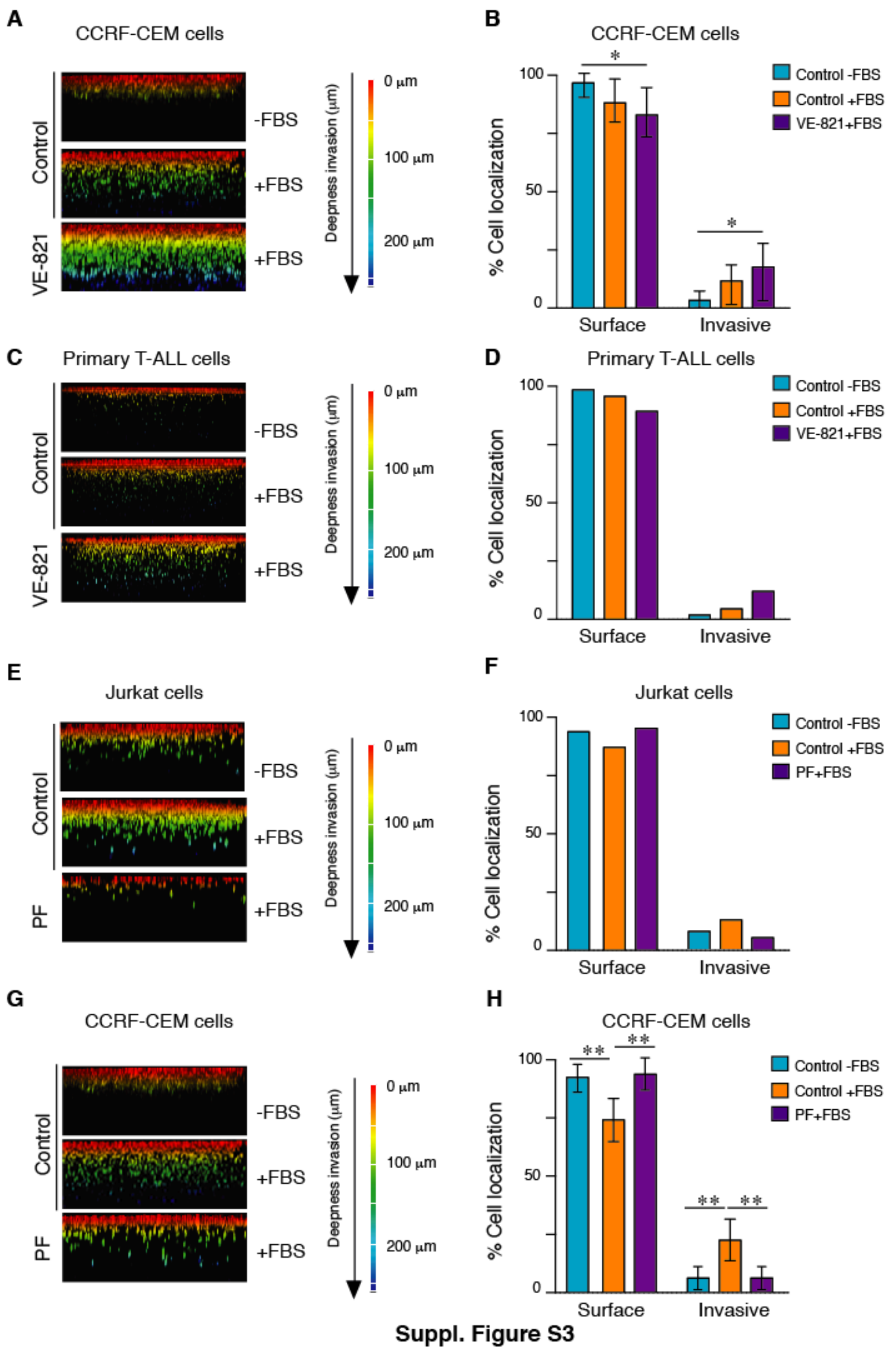


Suppl. Figure S3

**Supplementary Figure S3. ATR inhibition enhances migration through collagen I matrices in different T-ALL cell models. (A)** Representative 3D reconstructions of control and VE-821-treated CCRF-CEM cells migrating towards FBS through a 1.7 mg/mL collagen I matrix for 24 h. Cells were fixed and stained with Propidium Iodide. Colours indicate invasion depth (scale bar: 250 µm). **(B)** Quantification of the percentage of total cell-occupied area in the collagen gel. The upper third area was designated as cells at the surface, while cells in the lower two-thirds inside the collagen were designated as invasive cells ($N = 3$). * P<0.05, ** P<0.01. **(C)** Representative 3D reconstructions of control and VE-821-treated primary T-ALL cells migrating through a collagen I matrix for 48 h. Colours indicate invasion depth (scale bar: 250 μm). **(D)** Quantification of the percentage of total cell-occupied area in the gel. We designated the upper third as "Surface cells" (non-invasive), while cells in the lower two-thirds within the collagen were designated as "Invasive cells" **(E)** Representative 3D reconstructions of control and Chk-1-inhibited (250 nM PF-477736 for 24 h) Jurkat cells migrating through a collagen I matrix. Cells were fixed and stained with Propidium Iodide. Colours indicate invasion depth (scale bar: 250 μm). **(F)** Quantification of the percentage of total cell-occupied area located in the collagen gel. **(G)** Representative 3D reconstructions of control and Chk-1-inhibited CCRF-CEM cells migrating through a collagen I matrix for 24 h. Cells were fixed and stained with Propidium Iodide. Colours indicate invasion depth (scale bar: 250 μm). **(H)** Quantification of the percentage of total cell-occupied area in the collagen gel. ** P<0.01.

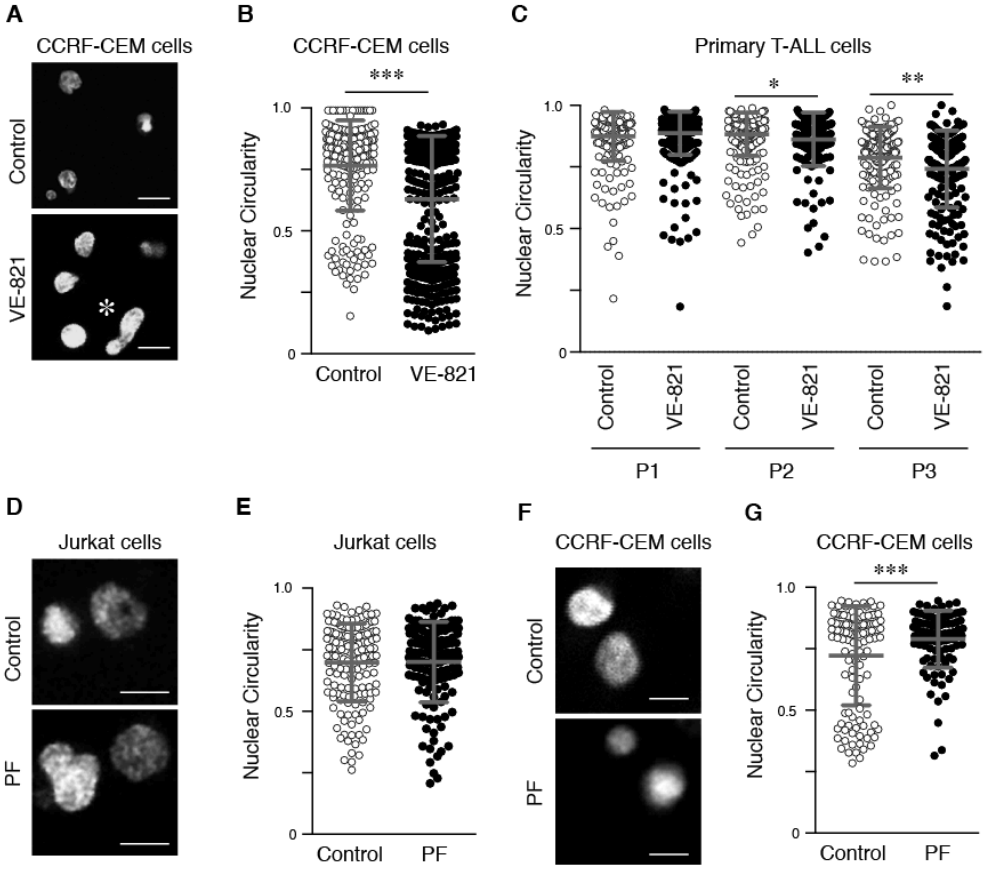


**Suppl. Figure S4**

**Supplementary Figure S4. Targeting ATR enhances nuclear deformation T-ALL cells through confined 3D environments. (A)** Representative immunofluorescence images of control and VE-821-treated CCRF-CEM cells embedded in a 1.7 mg/mL collagen I matrix for 4 h, fixed and stained with Hoechst (scale bar: 10 µm). **(B)** Nuclear circularity index quantification ($n$ = 236-403 cells; $N$ = 3). *** P<0.001. **(C)** Nuclear circularity index quantification of control and VE-821-treated primary T-ALL cells from three different patients ($n$ = 152-392 cells). * P<0.05, ** P<0.01. **(D)** Representative immunofluorescence images of control and Chk-1-inhibited Jurkat cells embedded in a collagen I matrix, fixed and stained with Hoechst (scale bar: 10 µm). **(E)** Nuclear circularity index quantification ($n$ = 155-137 nuclei; $N$ = 2). **(F)** Representative immunofluorescence images of control and Chk-1-inhibited CCRF-CEM cells embedded in a collagen I matrix, fixed and stained (scale bar: 10 µm). **(G)** Nuclear circularity index quantification ($n$ = 127-109 nuclei; $N$ = 2).

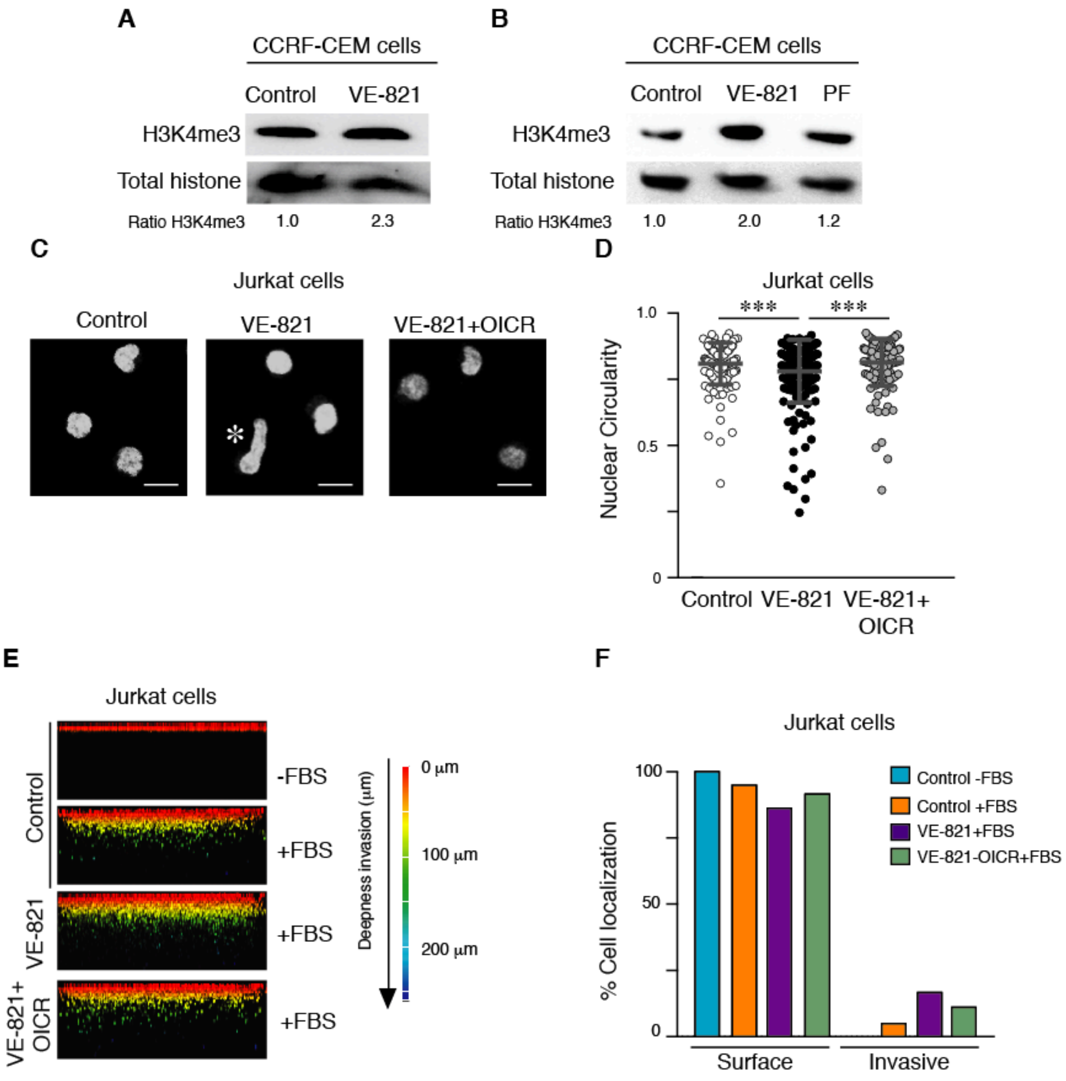


**Suppl. Figure S5**

**Supplementary Figure S5. ATR interference increases H3K4me3 levels and promotes WDR5-dependent nuclear deformation and invasion in T-ALL cells. (A)** Representative immunoblots showing H3K4me3 levels in total lysates from control and VE-821-treated CCRF-CEM cells. **(B)** Representative immunoblots showing H3K4me3 levels in total lysates from control, ATR-inhibited and Chk-1-inhibited Jurkat cells. **(C)** Jurkat cells were pretreated with OICR-9429 (1 μM) for 1 h before VE-821 addition. Then, cells were embedded in a 1.7 mg/mL collagen I matrix for 4 h, fixed, and stained with Hoechst (scale bar: 10 μm). **(D)** Nuclear circularity index quantification (*n* = 149-189 nuclei; *N* = 2). * P<0.05. **(E)** Representative 3D reconstructions of control, VE-821-treated, OICR-treated, and combined VE-821/OICR-treated Jurkat cells migrating through a collagen I matrix for 24 h. Cells were fixed and stained with Propidium Iodide. Colours indicate invasion depth (scale bar: 250 μm). **(F)** Quantification of the percentage of total cell-occupied area in the collagen gel. ($N = 3$).

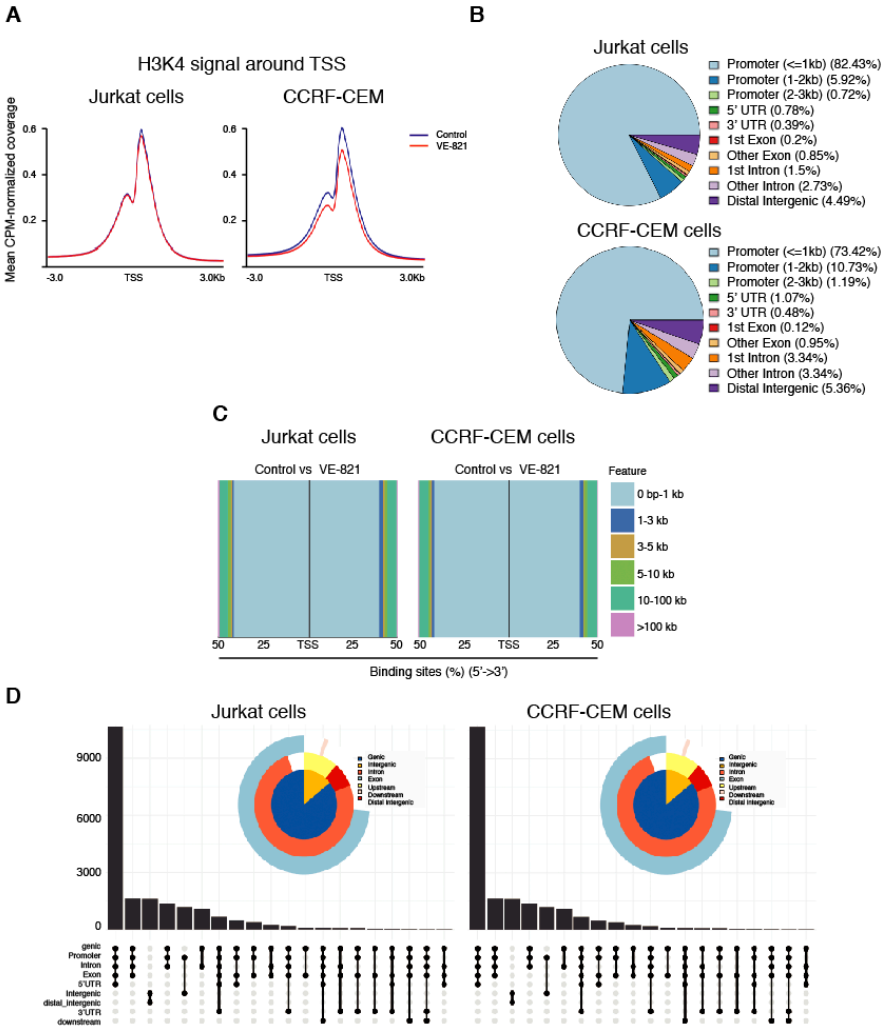


**Supplementary Figure S6**

**Supplementary Figure S6. Integrated ChIP-seq and RNA-seq analyses reveal chromatin and transcriptional changes following ATR inhibition in T-ALL cells. (A)** Line plots displaying the Mean CPM (counts per million)-normalized coverage of the H3K4me3 epigenetic signal centered around the Transcription Start Sites (TSS; ±3.0 kb) in Jurkat cells (left) and CCRF-CEM cells (right). The profiles compare control conditions (blue lines) against VE-821 treatment (red lines). **(B)** Pie charts showing the percentage distribution of H3K4me3 binding sites across distinct genomic features (including promoters with distance thresholds, UTRs, exons, introns, and distal intergenic regions) for Jurkat cells (top) and CCRF-CEM cells (bottom). **(C)** Bar charts depicting the relative percentage and physical distance (ranging from 0 bp to >100 kb) of genomic Features relative to the TSS (5' →

3') under control vs. VE-821 treatment conditions in both cell lines. **(D)** Combined bar charts and dot matrices (Upset plots) showing the multi-intersection frequency of genomic annotations, complemented by nested donut charts detailing the proportion of genic, promoter, intronic, exonic, UTR, upstream, downstream, and distal intergenic compartments for H3K4me3 peaks in Jurkat (left) and CCRF-CEM (right) cells.

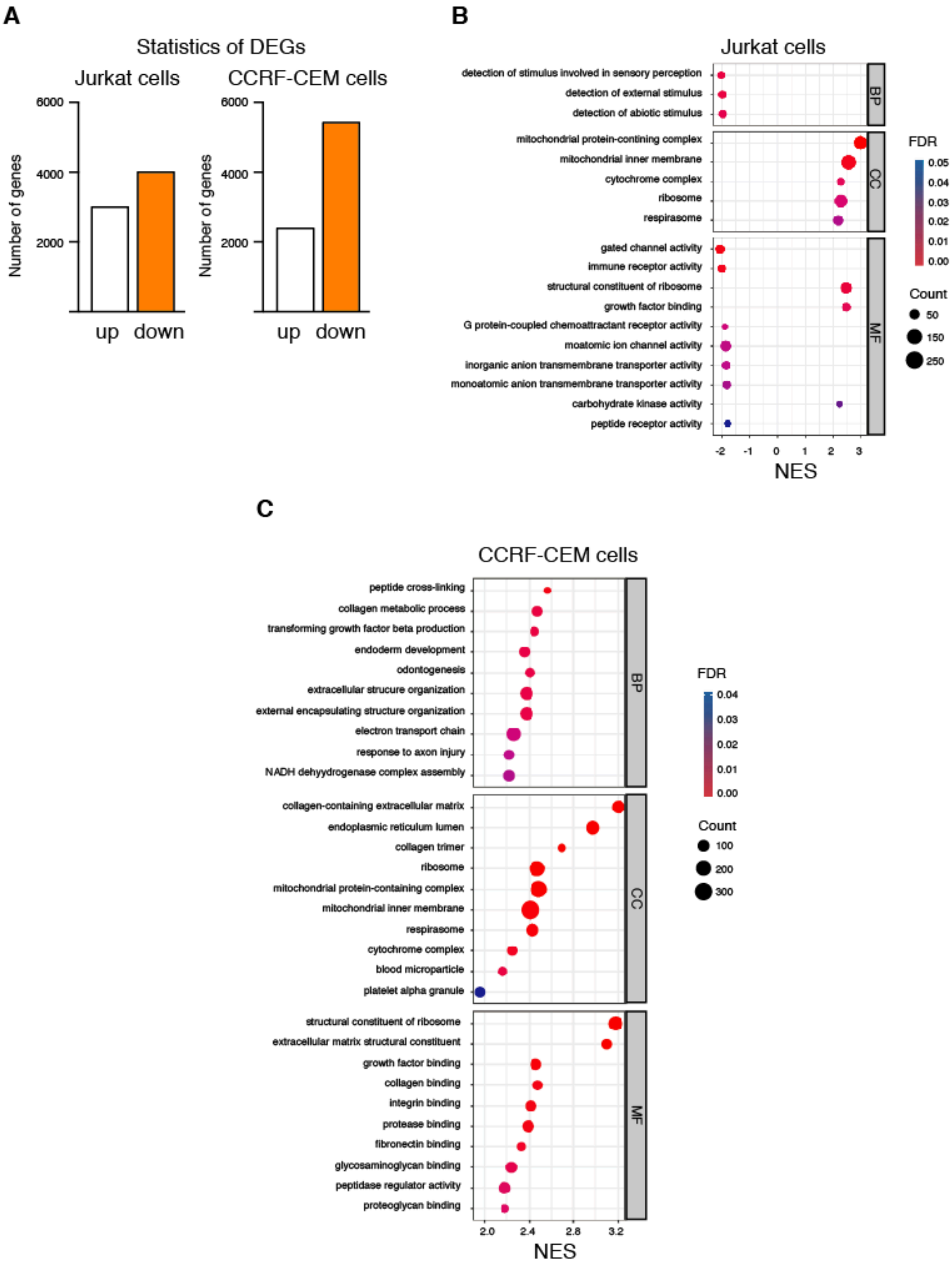


**Supl. Figure S7**

**Supplementary Figure S7. ATR inhibition induces cell-line-specific transcriptional and pathway enrichment responses in Jurkat and CCRF-CEM cells. (A)** Bar charts illustrating the total number of differentially expressed genes (DEGs) identified by RNA-seq that were significantly upregulated ("up") or downregulated ("down") following VE-821 treatment compared to control in Jurkat cells (left) and CCRF-CEM cells (right). **(B)** The graph shows the GSEA dot plot for Jurkat cells. Overrepresentation and enrichment analysis of functionally regulated pathways. The x-axis represents the Normalized Enrichment Score

(NES). The size of the dots indicates the gene Count involved in each term, and the color gradient reflects the False Discovery Rate (FDR) statistical significance. Right labels denote the GO categories: Biological Process (BP), Cellular Component (CC), and Molecular Function (MF). **(C)** GSEA dot plot for CCRF-CEM cells: Enriched functional categories (BP, CC, and MF) in CCRF-CEM cells under VE-821 treatment. Dot distribution is mapped against the NES value on the x-axis, with dot size representing the gene Count and color corresponding to the FDR value.

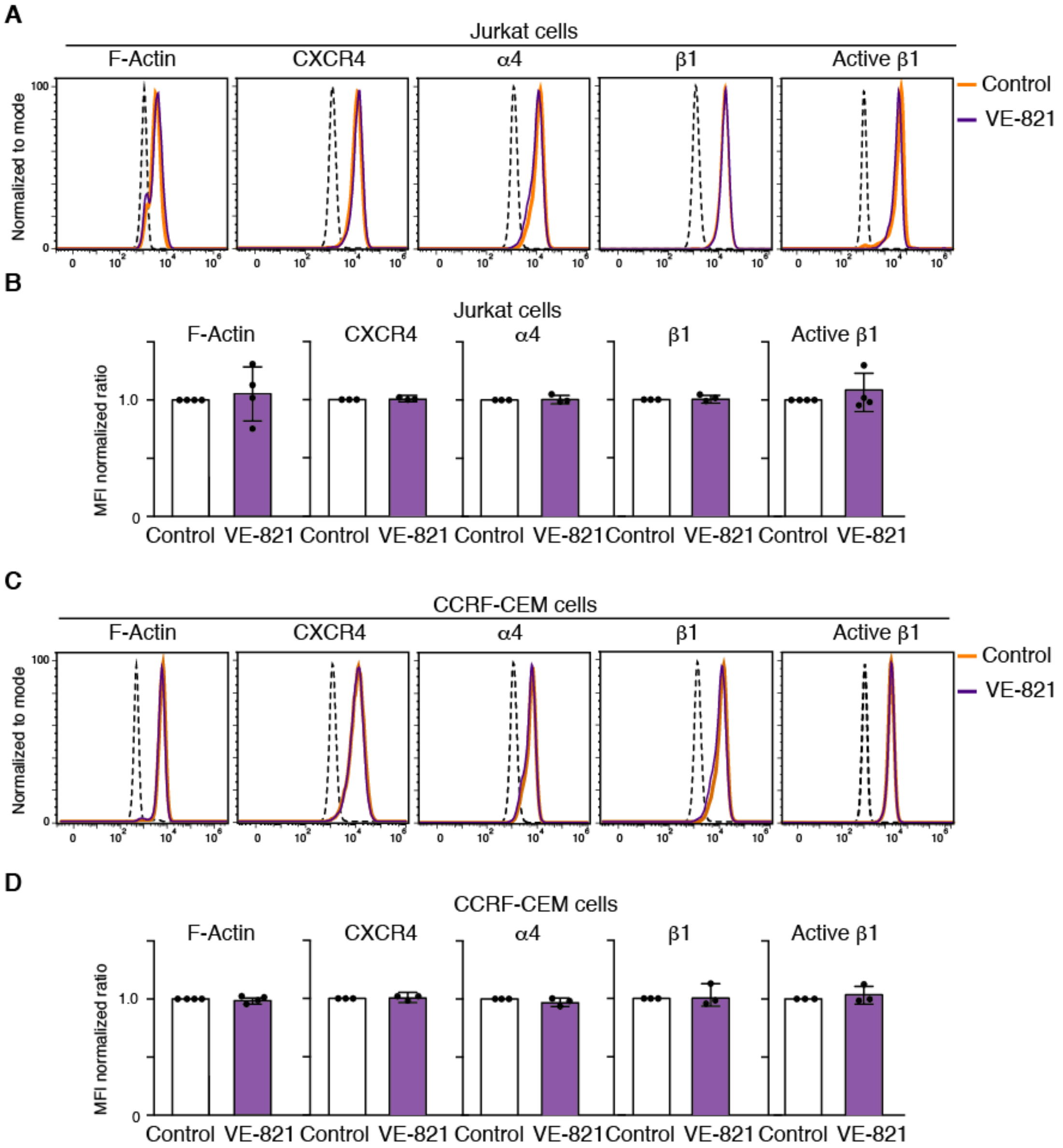


**Supl. Figure S8**

**Supplementary Figure S8. ATR inhibition maintains the expression of cytoskeletal and adhesion-associated markers. (A)** Flow cytometry analysis of F-actin, CXCR4, integrin α4, integrin β1, and active integrin β1 expression in control and VE-821-treated Jurkat cells. **(B)** Quantification of the normalized mean fluorescence intensity (MFI) ratio for F-actin, CXCR4, integrin α4, integrin β1, and active integrin β1 in control and VE-821-treated Jurkat cells. **(C)** Flow cytometry analysis of F-actin, CXCR4, integrin α4, integrin β1, and active integrin β1 expression in control and VE-821-treated CCRF-CEM cells. **(D)** Quantification of the normalized MFI ratio for F-actin, CXCR4, integrin α4, integrin β1, and active integrin β1 in control and VE-821-treated CCRF-CEM cells.

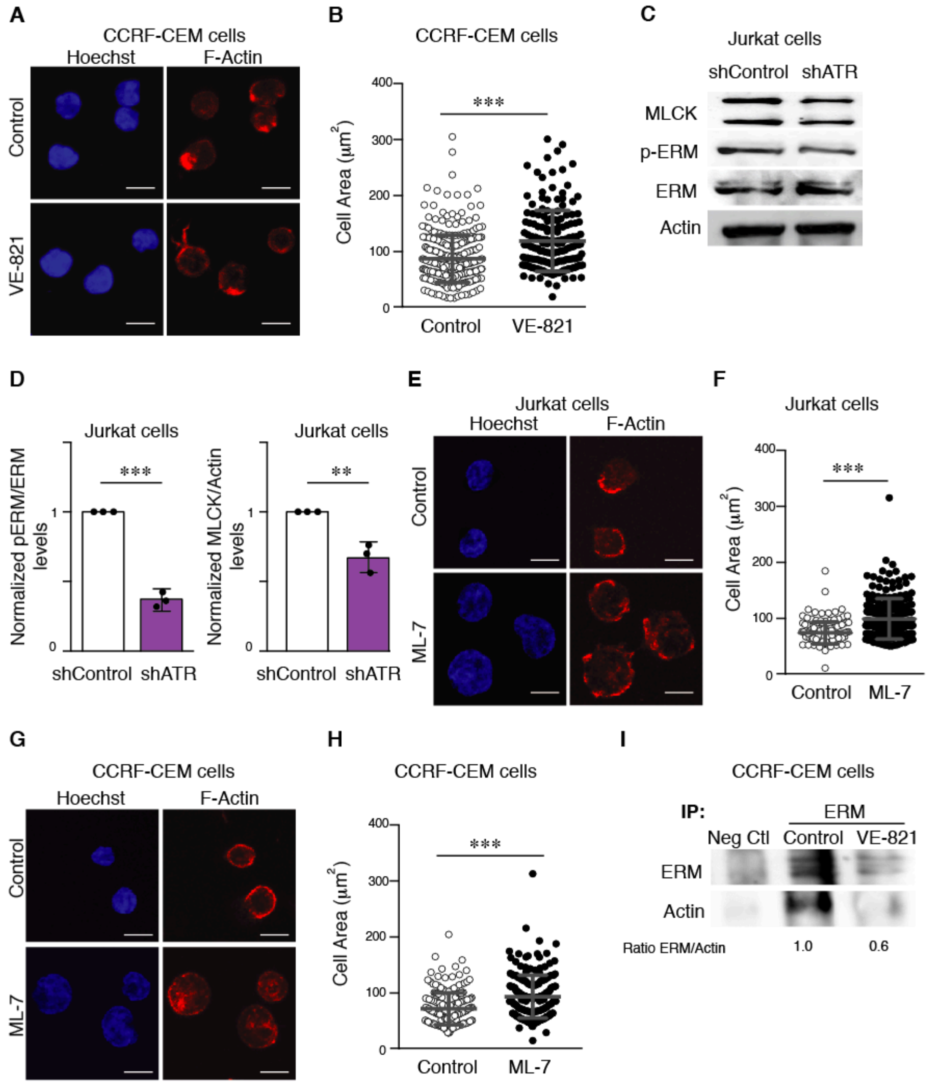


**Supl. Figure S9**

**Supplementary Figure S9. ATR inhibition alters cell adhesion, cell spreading, and cortical cytoskeleton regulation in T-ALL cells. (A)** Representative immunofluorescence images of control and VE-821-treated CCRF-CEM cells cultured on poly-L-lysine-coated slides for 1 h, fixed, and stained with phalloidin (red) and Hoechst (blue) (scale bar: 10 µm). **(B)** Cellular area quantification from phalloidin-staining ($n$ = 328-171 cells; $N$ = 3). *** P<0.001. **(C)** Representative immunoblots for MLCK, ERM and phospho-ERM proteins in whole-cell lysates of control (shControl) and ATR-depleted (shATR) Jurkat cells. **(D)** phospho-ERM/ERM and MLCK/Actin ratios. ($N$ = 3). **(E)** Representative immunofluorescence images of control (untreated) and MLCK-inhibited (20 µM ML-7 for 20 min) Jurkat cells cultured on poly-L-lysine-coated slides, fixed, and stained with phalloidin (red) and Hoechst (blue) (scale bar: 10 μm). **(F)** Cellular area quantification (*n* = 142-206 cells). *** $P<0.001$. **(G)** Representative immunofluorescence images of control (untreated) and MLCK-inhibited (20 μM ML-7 for 20 min) CCRF-CEM cells cultured on poly-L-lysine-coated slides and stained as in (E) (blue) (scale bar: 10 μm). **(H)** Cellular area quantification (*n* = 222-214 cells). *** $P<0.001$. **(I)** Representative immunoblots showing actin–ERM interactions following immunoprecipitation of ERM from control and VE-821-treated CCRF-CEM cell lysates. Actin was detected by immunoblotting in ERM immunoprecipitates. Neg. Ctl. represents the negative control immunoprecipitation. Numbers indicate the protein quantification ratio after normalizing control cell values to 1.
