## Supplementary material for "ATR coordinates nuclear and cytoskeletal mechanics during leukemic migration": Table 1

| Characteristic | P1 | P2 | P3 | P4 |
|---|---|---|---|---|
| **Gender** | Male | Female | Female | Male |
| **Risk group** | Intermediate | High | Intermediate | High |
| **WBC 1 (>50,000 cells/mm$^3$)** | 367,000 | 350,000 | 32,800 | 16,100 |
| **MRD 1 after induction** | < 0,01 | 0.23 | < 0,01 | 1.16 |
| **ETV6/RUNX1** | 46,XY, t(11;14)(p13;q11.2) LMO2-TRD, t(7;8)(q34,q24.21) with deletion TRB (7q34), del(9)(p21.3) CDKN2A/B, del(4)(q25) LEF1 | 46,XX, del(1)(p33), del(10)(q23.31) | No | 49,XY, +8, +10, +19 |
| **MLL rearrangements** | — | — | No | — |
| **Hyperdiploidy** | No | No | No | Yes |
| **Relapse** | No | No | No | No |
| **Death** | No | No | No | No |
