## Supplementary material for "ATR coordinates nuclear and cytoskeletal mechanics during leukemic migration": Table 2

Jurkat_ChIPseq-vs-RNSseq

| ID | gene_symbol | logFC_ChIPseq H3K4me3 CEM | Pvalue_ChIPseq H3K4me3 CEM | logFC_RNAseq CEM | Pvalue_RNAseq CEM | sig | direction | signficance |
|---|---|---|---|---|---|---|---|---|
| **ENSG00000125170** | DOK4 | -0.980252769 | 0.039988592 | -0.454571574 | 8.35E-06 | sig | both_down | sig_both_down |
| **ENSG00000133243** | BTBD2 | -0.522195216 | 0.039248138 | 0.295376728 | 2.91E-21 | sig | opposite | sig_opposite |
| **ENSG00000140470** | ADAMTS17 | -3.843989484 | 0.033369552 | -1.509408754 | 1.55E-11 | sig | both_down | sig_both_down |
| **ENSG00000147488** | ST18 | -3.843989484 | 0.033369552 | 3.955259514 | 0.006111436 | sig | opposite | sig_opposite |
| **ENSG00000167984** | NLRC3 | -2.586033337 | 0.03860956 | -0.37096161 | 3.15E-21 | sig | both_down | sig_both_down |

CCRF-CEM_ChIPseq-vs-RNSseq

| ID | gene_symbol | logFC_ChIPseq H3K4me3 CEM | Pvalue_ChIPseq H3K4me3 CEM | logFC_RNAseq CEM | Pvalue_RNAseq CEM | sig | direction | signficance |
|---|---|---|---|---|---|---|---|---|
| **ENSG00000014123** | UFL1 | -0.352200408 | 0.020769114 | -0.817172288 | 6.72E-66 | sig | both_down | sig_both_down |
| **ENSG00000066651** | TRMT11 | -0.405899571 | 0.035252808 | -0.348128921 | 7.61E-06 | sig | both_down | sig_both_down |
| **ENSG00000072803** | FBXW11 | 3.506900393 | 0.00815548 | -0.384802829 | 8.47E-16 | sig | opposite | sig_opposite |
| **ENSG00000103489** | XYLT1 | 1.486766652 | 0.037121731 | -0.532174101 | 6.23E-32 | sig | opposite | sig_opposite |
| **ENSG00000113360** | DROSHA | 0.257711387 | 0.035127246 | -0.162006661 | 2.81E-07 | sig | opposite | sig_opposite |
| **ENSG00000132394** | EEFSEC | 4.282657718 | 0.039509657 | 0.273063472 | 2.20E-09 | sig | both_up | sig_both_up |
| **ENSG00000133247** | KMT5C | -0.297006257 | 0.049507365 | 0.214209222 | 0.000190102 | sig | opposite | sig_opposite |
| **ENSG00000141337** | ARSG | 2.895869205 | 0.046251106 | -0.316431528 | 0.001976679 | sig | opposite | sig_opposite |
| **ENSG00000156398** | SFXN2 | -0.291241354 | 0.036467439 | -0.157623232 | 0.009118109 | sig | both_down | sig_both_down |
| **ENSG00000159917** | ZNF235 | 0.344452612 | 0.042507217 | -0.432418242 | 0.003980711 | sig | opposite | sig_opposite |
| **ENSG00000160219** | GAB3 | 3.091292106 | 0.013786663 | -0.677848199 | 0.002891706 | sig | opposite | sig_opposite |
| **ENSG00000160360** | GPSM1 | 1.68395382 | 0.029102685 | 0.470414137 | 0.000811679 | sig | both_up | sig_both_up |
| **ENSG00000162783** | IER5 | 0.686164061 | 0.036675263 | 0.252726265 | 1.12E-07 | sig | both_up | sig_both_up |
| **ENSG00000163541** | SUCLG1 | -0.319900652 | 0.021892114 | 0.170494632 | 1.09E-08 | sig | opposite | sig_opposite |
| **ENSG00000174255** | ZNF80 | 1.18651256 | 0.030979504 | -0.852526731 | 0.006319287 | sig | opposite | sig_opposite |
| **ENSG00000178105** | DDX10 | 3.869839804 | 0.015812818 | 0.180359889 | 1.14E-05 | sig | both_up | sig_both_up |
| **ENSG00000254858** | MPV17L2 | 1.512235523 | 0.038559502 | 0.330928584 | 1.15E-09 | sig | both_up | sig_both_up |
